# The Ets transcription factor ETV4 regulates FGF1-dependent proliferation and glycolysis in ER-positive breast cancer

**DOI:** 10.64898/2026.01.13.699240

**Authors:** Barbara Mensah Sankofi, Nisha S. Thomas, Malika Sekhri, Haoning Howard Cen, William L. Berry, Stevi J. Murguia, Rameswari Velayutham, Michael C. Rudolph, Elizabeth A. Wellberg

## Abstract

Obesity is a risk factor for estrogen receptor (ER) positive breast cancer. Beyond body mass index, adult weight gain increases breast cancer risk. During weight gain, hypertrophic adipocytes produce fibroblast growth factor 1 (FGF1), which drives estrogen-independent growth of ER-positive tumors. Effects of FGF1 on breast cancer cells include elevated proliferation and enhanced glycolytic activity. We identified the Ets transcription factor ETV4 as a target of FGF1 treatment across multiple breast cancer cell lines. Our objective was to define the role of ETV4 in mediating the tumor-promotional effects of FGF1, to better understand how weight gain and obesity drive breast cancer risk and progression. Here, we determined that ETV4 directly associates with a poor prognosis for patients with ER-positive tumors and positively correlates with FGF1 levels in the context of obesity. We demonstrate that ETV4 is required to mediate the pro-tumorigenic effects of FGF1 on cell proliferation, glycolytic reprogramming, and tamoxifen sensitivity *in vitro*, and on tumor growth in the presence of estrogen in obese mice. *In vitro*, ETV4 overexpression enhances proliferation and metabolic activity, mimicking effects of FGF1 on breast cancer cells, but it is not sufficient to promote ER-positive tumor growth before or after estrogen deprivation *in vivo* in lean females. This study reveals a potentially novel mechanism through which weight gain, characterized by excess FGF1 production, drives the development of aggressive features in the prevalent ER-positive breast cancer subtype.

## INTRODUCTION

Breast cancer remains the most diagnosed malignancy and the leading cause of cancer-related death among women globally [1]. Estrogen receptor (ER)-positive tumors comprise 70% of all cases and are treated with endocrine therapies that antagonize ER activity or deplete estrogen production. Although endocrine therapies have substantially improved prognosis, some patients develop resistance after treatment, leading to cancer recurrence and metastatic disease [2, 3]. In recent years, the growing prevalence of obesity has increased the risk for and mortality from breast cancer [4–6]. Compared with ER-negative tumors, those expressing ER are promoted by obesity, particularly after menopause [6–9].

Obesity is associated with elevated estrogen levels, which likely explains its relationship with breast cancer incidence [10]. However, aromatase inhibitors that block estrogen production significantly reduce circulating estrogens even in women with a high BMI [11, 12], suggesting that obesity-associated ER-positive tumor progression after treatment is driven by factors other than excess estrogen. In preclinical models, we found that adult weight gain promotes the growth of ER-positive tumors after estrogen withdrawal (EWD) treatment through adipose-derived fibroblast growth factor 1 (FGF1) [13, 14]. Adipose tissue is abundant in the breast microenvironment and hypertrophic adipocytes are a source of FGF1, especially during weight gain [14, 15], which is accelerated by the loss of estrogen. In mammary adipose tissue, FGF1 expression correlated with obesity and was upregulated during weight gain in mice, rats, and humans [13, 14]. In addition, breast cancer patient-derived xenograft (PDX) and primary human tumors had elevated levels of phosphorylated fibroblast growth factor receptor 1 (FGFR1) in the context of obesity [13]. Inhibition of FGFR signaling *in vivo* restored sensitivity of tumors to EWD in obese females, highlighting a potential mechanism connecting weight gain to breast cancer progression.

In three ER-positive breast cancer cell lines, we found that FGF1 treatment enhanced glycolytic metabolism and upregulated expression of the Ets transcription factor, ETV4 [16]. ETV4 has diverse cellular functions, including during embryogenesis [17, 18]. In the developing mammary bud, ETV4 is expressed throughout the luminal epithelial cell population, and it is required for appropriate branching morphogenesis of the mammary ductal network during puberty and pregnancy [19]. ETV4 is upregulated in breast tumor compared with normal tissue and associates with a relatively poor patient prognosis [20, 21]; however, much of the published work has focused on HER2-positive or triple negative breast cancer, where ETV4 is demonstrated to have tumor promotional effects [22, 23]. During mammalian development, FGF-FGFR signaling through ETV4 is implicated in limb patterning, in the formation of lachrymal glands and lenses [24–26], and in the progression of synovial sarcoma [27]. In breast cancer cells, one study showed that FGF2 induced ETV4 expression in MDA-231 cells and promoted binding to Ets-response elements in MCF7 cells [28]. In endometrial cancer, ETV4 regulates ER transcriptional activity [29]; however, FGFR signaling has not been implicated ETV4 action in ER-positive breast cancer.

In this study, we aimed to further define the mechanisms linking obesity to breast cancer by determining how ETV4 mediates the effects of FGF1 on ER-positive cancer cells, emphasizing proliferation, glycolytic activity, gene expression, and tumor growth *in vivo*. We show that high ETV4 expression in tumors is positively associated with poor prognosis for patients with ER-positive breast cancer, and that ETV4 is necessary to mediate the tumor promotional effects of FGF1 *in vitro* and *in vivo*. We also demonstrate that, while ETV4 is sufficient to elicit similar pro-tumorigenic effects as FGF1 on cancer cells *in vitro*, it is not capable of supporting estrogen-independent tumor growth *in vivo*, suggesting that additional physiologic mechanisms mediate resistance to endocrine therapy.

## RESULTS

### ETV4 associates with obesity, FGF1, and poor prognosis in ER-positive breast cancer

We previously reported that ETV4 was induced by FGF1 treatment of some ER-positive breast cancer cells *in vitro* [16]. To investigate the role of ETV4 in breast cancer progression, we first looked at human breast tissues using publicly available transcriptomic data [30]. ETV4 levels were higher in invasive breast tumors compared to matched normal samples (Fig 1A). With all subtypes combined, high tumor ETV4 expression associated with a shorter overall survival for breast cancer patients (Fig 1B), as has been demonstrated previously [22, 23]. When comparing tumors based on ER status, we found that high ETV4 expression (mRNA z-score >1.5) was overrepresented in ER-negative versus ER-positive tumors (Fig 1C). ER-negative breast cancer associates with a relatively low survival, which could explain why high ETV4 expression predicts a poor outcome. When we specifically evaluated ER-positive tumors, high ETV4 expression also associated with shorter overall (Fig 1D), and recurrence-free survival intervals (Fig 1E). Next, we investigated the relationship between ETV4 expression and pathologic complete response after cancer treatment [31]. Focusing only on ER-positive tumors, ETV4 levels were higher in those that were non-responsive to aromatase inhibitor therapy (Fig 1F) or to any chemotherapy (Fig 1G), compared with responsive tumors. Using the same datasets, we evaluated the utility of ETV4 as a predictive biomarker. Receiver operating characteristic (ROC) curve analysis showed that, although statistically significant, the area under the curve was less than 0.7 for ETV4 expression in ER-positive tumors treated with aromatase inhibitors (Supplemental Fig 1A) or any chemotherapy (Supplemental Fig 1B), suggesting limited utility as a prognostic biomarker on its own [32]. Together, these data indicate that although ETV4 expression is higher in tumors that are non-responsive to therapy, additional factors contribute to whether ETV4 alone is a reliable prognostic biomarker.

**Figure 1.**
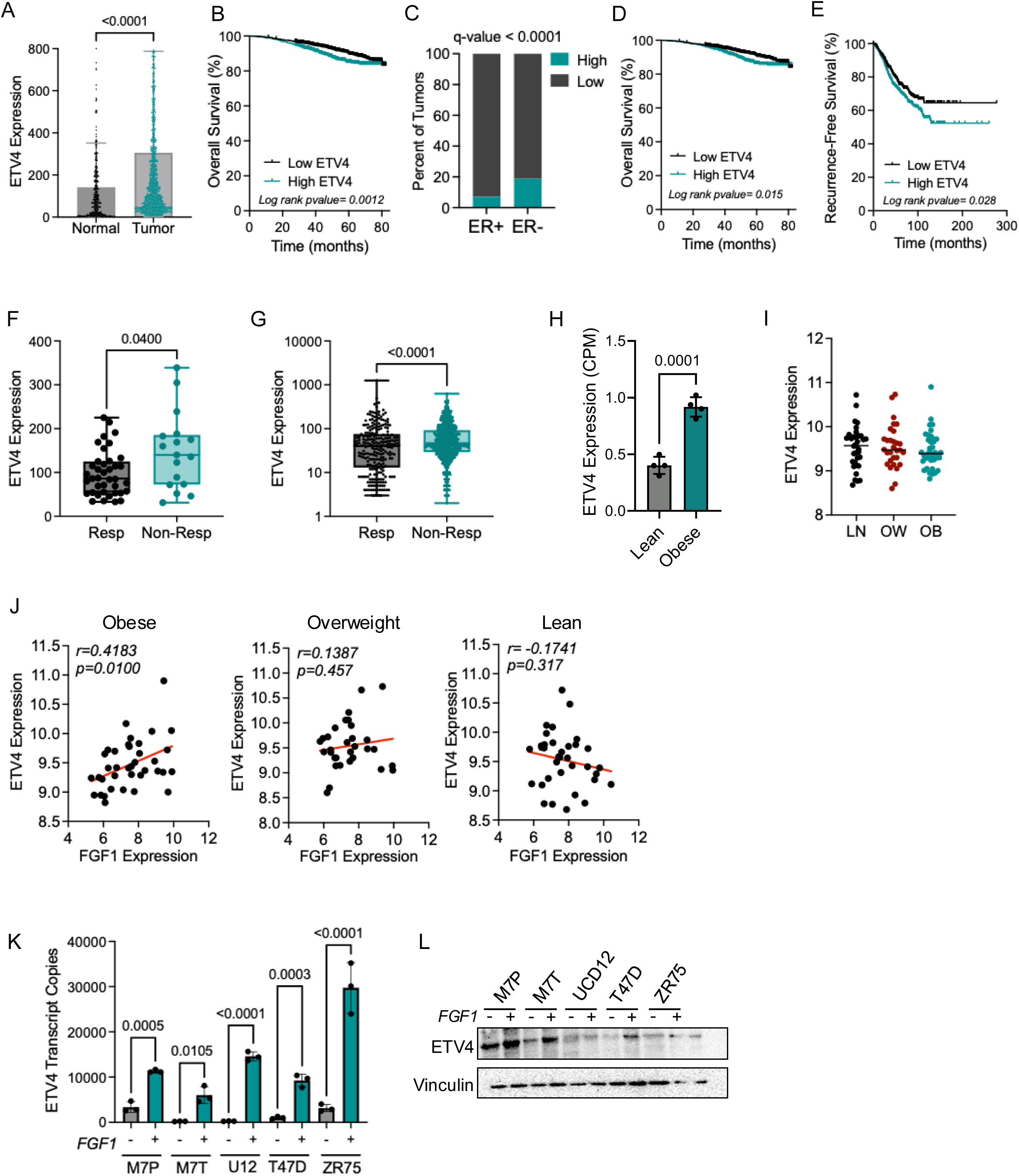
High ETV4 expression associates with patient survival and FGF1 expression. (A) ETV4 expression in breast tumors and paired normal breast tissue from people with breast cancer. Data obtained from tnmplot.com. Mann-Whitney test determined significance. (B) High ETV4 expression (RNA seq) associates with lower overall survival for people with breast cancer. HR=1.44 (1.14-1.82). (C) Tumors with high ETV4 expression (mRNA z-score >1.5) are more likely to be ER-negative than ER-positive. Chi-squared test q-value <0.0001. (D) High ETV4 expression (RNA seq) associates with lower overall survival for patients with ER-positive breast cancer. HR=1.39 (1.05-1.85). (E) High ETV4 expression (array) associates with lower recurrence-free survival for patients with lymph node-positive ER-positive breast cancer. HR=1.3 (1.031-1.628). (F) ETV4 expression in ER-positive breast tumors stratified by pathologic complete response after aromatase inhibitors using data from ROCplotter.com. (G) ETV4 expression in ER-positive tumors stratified by pathologic complete response to any chemotherapy using data from ROCplotter.com. Mann-Whitney test determined significance for *f* and *g*. (H) ETV4 expression (RNA seq) in UCD12 PDX tumors from lean or obese female mice. (I) ETV4 expression (array) in tumors from patients with ER-positive breast cancer (data from GSE24185). (J) Pearson correlation between ETV4 and FGF1 expression in tumors from patients classified as obese, overweight, or lean based on BMI (data from GSE24185). (K) Expression of ETV4 in ER-positive breast cancer cells with or without FGF1 treatment. MCF7 Parental (M7P) or TAMR (M7T); UCD12 (U12). (L) Representative western blot showing ETV4 expression in ER-positive breast cancer cells with or without FGF1 treatment.

Our earlier work found that obesity supports estrogen-independent growth of ER-positive tumors, associated with mammary adipose production of FGF1 during weight gain and tumor FGFR activation [13]. To determine the relationship between ETV4 expression, obesity, and FGF1, we evaluated tumors from mice and publicly available data from breast cancer patients. ETV4 expression was greater in ER-positive breast cancer PDX tumors grown in obese compared with lean female mice (Fig 1H) [13]. However, in primary human ER-positive breast tumors, ETV4 was similarly expressed across lean, overweight, and obese BMI categories (Fig 1I). Based on our work demonstrating excess FGF1 production in the obese tumor microenvironment [13], we asked how ETV4 and FGF1 expression were related in human tumors across BMI categories. ETV4 and FGF1 were directly and significantly correlated in tumors from patients with obesity (Fig 1J), but not in those patients who were classified as overweight or lean. In several ER-positive breast cancer cell lines, FGF1 treatment markedly induced ETV4 gene (Fig 1K) and protein expression (Fig 1L). The difference in obesity-associated ETV4 expression between tumors from mice and humans illustrates that, in humans, breast cancer is driven by complex mechanisms that cannot be explained by BMI alone. Collectively, these data show that ETV4 correlates strongly with FGF1 and is linked to poor prognosis and disease progression in ER-positive breast cancer.

### ETV4 mediates effects of FGF1 on cell proliferation

In breast cancer cells, we showed that FGF1 enhances proliferation and elevates glycolytic activity [16]. To investigate whether ETV4 is required to mediate these effects, we knocked down expression in three ER-positive breast cancer cell lines: parental MCF7 cells (MCF7-P; Fig 2A), tamoxifen-resistant MCF7 cells (MCF7-TAMR; Fig 2B), and UCD12 cells (Supplemental Fig 2A), which were derived from a breast cancer PDX [33]. These cell lines previously demonstrated differential dependence on estrogen in obese mice when grown as tumors *in vivo*; MCF7-TAMR and UCD12 tumors continued to grow after withdrawal of supplemental estrogen in obese females, while parental MCF7 (MCF7-P) did not [16]. In all three cell lines expressing shRNA control, FGF1 treatment increased proliferation compared to vehicle (Fig 2C-F; Supplemental Fig 2B-C), which is consistent with our prior studies [16]. In MCF7-P cells, knockdown of ETV4 did not impact proliferation at baseline, but attenuated cell growth after FGF1 treatment (Fig 2C-D). In contrast, MCF7-TAMR (Fig 2E-F) and UCD12 cells (Supplemental Fig 2B-C) lacking ETV4 had lower levels of proliferation, regardless of FGF1 treatment. Thus, while the requirement of ETV4 to sustain basal proliferation was different across cell lines, their response to FGF1 was consistently impaired when ETV4 was reduced.

**Figure 2.**
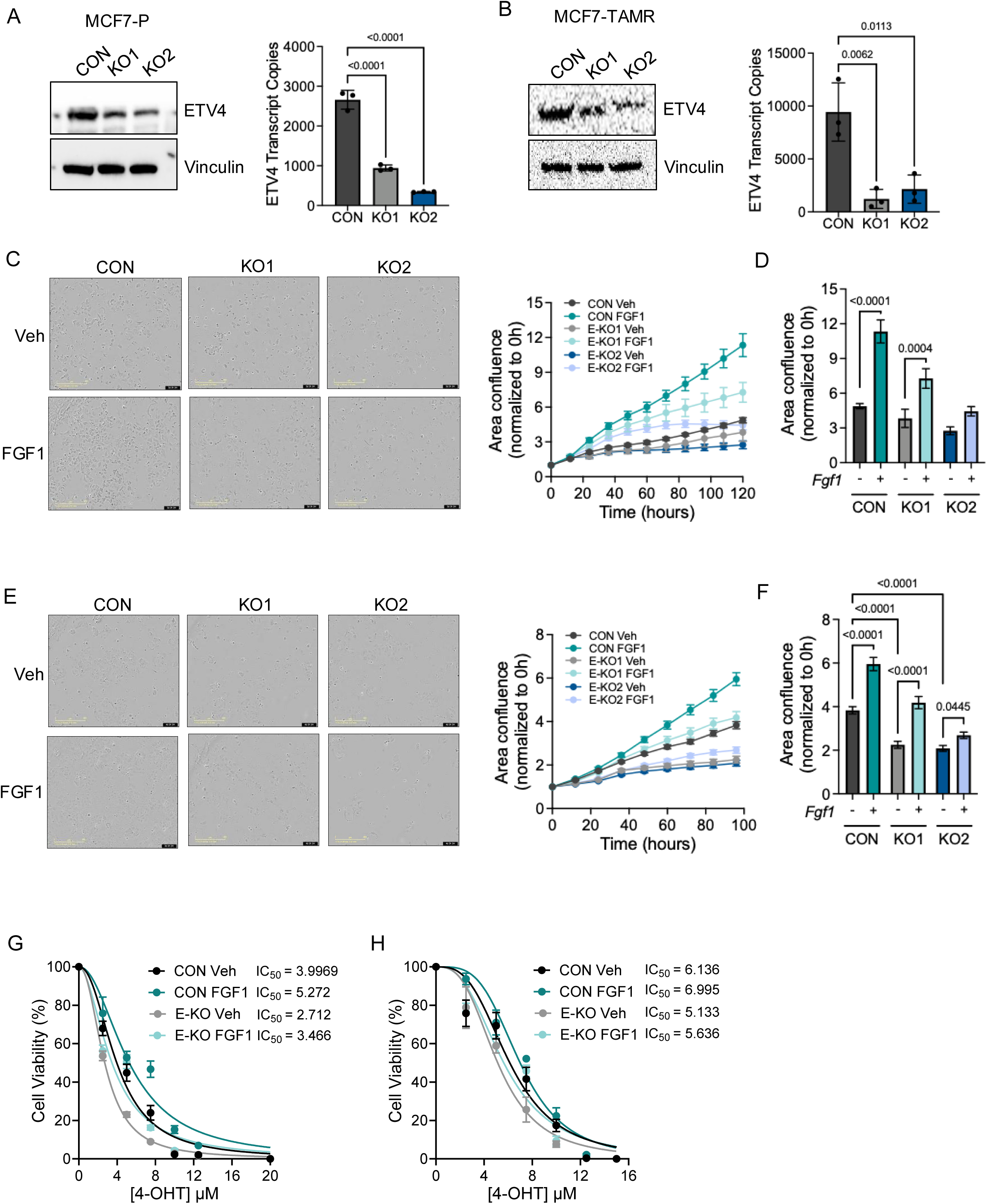
ETV4 is necessary for FGF1-mediated proliferation in ER-positive breast cancer cells. (A) Q-PCR (*left*) and immunoblot (*right*) analysis of ETV4 in MCF7-P cells. (B) Q-PCR (*left*) and immunoblot (*right*) analysis of ETV4 in MCF7-TAMR cells. (C) Representative images of the final timepoint (left) and growth rates (right) of MCF7-P control or ETV4-knockdown cells treated with vehicle or FGF1. (D) Area confluence relative to control vehicle time 0 of cells at the final timepoint following treatment of MCF7-P cells. Two-way ANOVA testing for main effects of ETV4 knockdown or FGF1 treatment or interaction was performed. P-values indicate post-hoc multiple testing for specific differences between pre-defined comparisons. (E) Representative images of the final timepoint (left) and growth rates (right) of MCF7-TAMR control or ETV4-knockdown cells treated with vehicle or FGF1. (F) Area confluence relative to control vehicle time 0 of cells at the final timepoint following treatment of MCF7-TAMR cells. Two-way ANOVA testing for main effects of ETV4 knockdown or FGF1 treatment or interaction was performed. P-values indicate post-hoc multiple testing for specific differences between pre-defined comparisons. (G) Dose-response curve and interpolated IC_50_ values of tamoxifen treatment in MCF7-P control and ETV4 knockdown cells following treatment with or without FGF1. (H) Dose-response curve and interpolated IC_50_ values of tamoxifen treatment in MCF7-TAMR control and ETV4 knockdown cells following treatment with or without FGF1.

Next, we asked how FGF1 and ETV4 influenced the sensitivity of breast cancer cells to tamoxifen, which is a standard therapy for ER-positive breast cancer. Treatment of MCF7-P cells with FGF1 lowered tamoxifen sensitivity, shifting the viability curve to the right (Fig 2G; Additional File 1). Knockdown of ETV4 had the opposite effect, increasing tamoxifen sensitivity in the absence of FGF1. Upon addition of FGF1, cells became slightly more resistant to tamoxifen, however FGF1 treatment of ETV4 knockdown cells was not sufficient to restore viability to that seen in control cells (Fig 2G; Additional File 1). MCF7-TAMR cells responded similarly, although the magnitude of effect was smaller than in MCF7-P cells. As expected, MCF7-TAMR control cells were less sensitive to tamoxifen compared with MCF7-P cells (Fig 2H; Additional File 1). FGF1 treatment promoted greater tamoxifen resistance in MCF7-TAMR controls, and knockdown of ETV4 sensitized cells to tamoxifen. Adding FGF1 to ETV4 knockdown cells had a minor effect on restoring tamoxifen resistance but did not reach the levels seen in control cells (Fig 2H; Additional File 1). Together, these data indicate that FGF1 supports tamoxifen resistance in breast cancer cells mediated in part through ETV4.

To determine whether ETV4 was sufficient to enhance cell proliferation, we generated ETV4-overexpressing MCF7-P (Fig 3A) and MCF7-TAMR (Fig 3B) cells. Treatment of control MCF7-P cells with FGF1 increased proliferation (Fig 3C-D), consistent with data from ETV4 knockdown experiments. Overexpression of ETV4 similarly promoted proliferation, and addition of FGF1 did not further increase this endpoint (Fig 3C-D). Proliferation of MCF7-TAMR control cells was also increased by FGF1 treatment and ETV4 overexpression (Fig 3E-F). Addition of FGF1 to ETV4-overexpressing cells further enhanced proliferation compared with vehicle control (Fig 3E-F). ETV4 overexpression in MCF7-P cells modestly increased tamoxifen resistance (Fig 3G; Additional File 1), and similar effects were seen in MCF7-TAMR cells (Fig 3H). To determine how FGFR activation was impacted by ETV4, we compared MCF7-P to MCF7-TAMR cells. Interestingly, MCF7-TAMR cells were significantly more resistant to FGFR inhibition (with BGJ398) compared to MCF7-P cells (Fig 3I; Additional File 1). Overexpression of ETV4 in MCF7-P cells also greatly reduced sensitivity to BGJ398 but had less of an effect on MCF7-TAMR cells (Fig 3I). Together, these data indicate that ETV4 is necessary to mediate the robust proliferation of ER-positive breast cancer cells in response to FGF1 and is sufficient to stimulate proliferation on its own. Furthermore, ETV4 plays a role in the sensitivity of breast cancer cells to tamoxifen and to pharmacologic FGFR inhibition.

**Figure 3.**
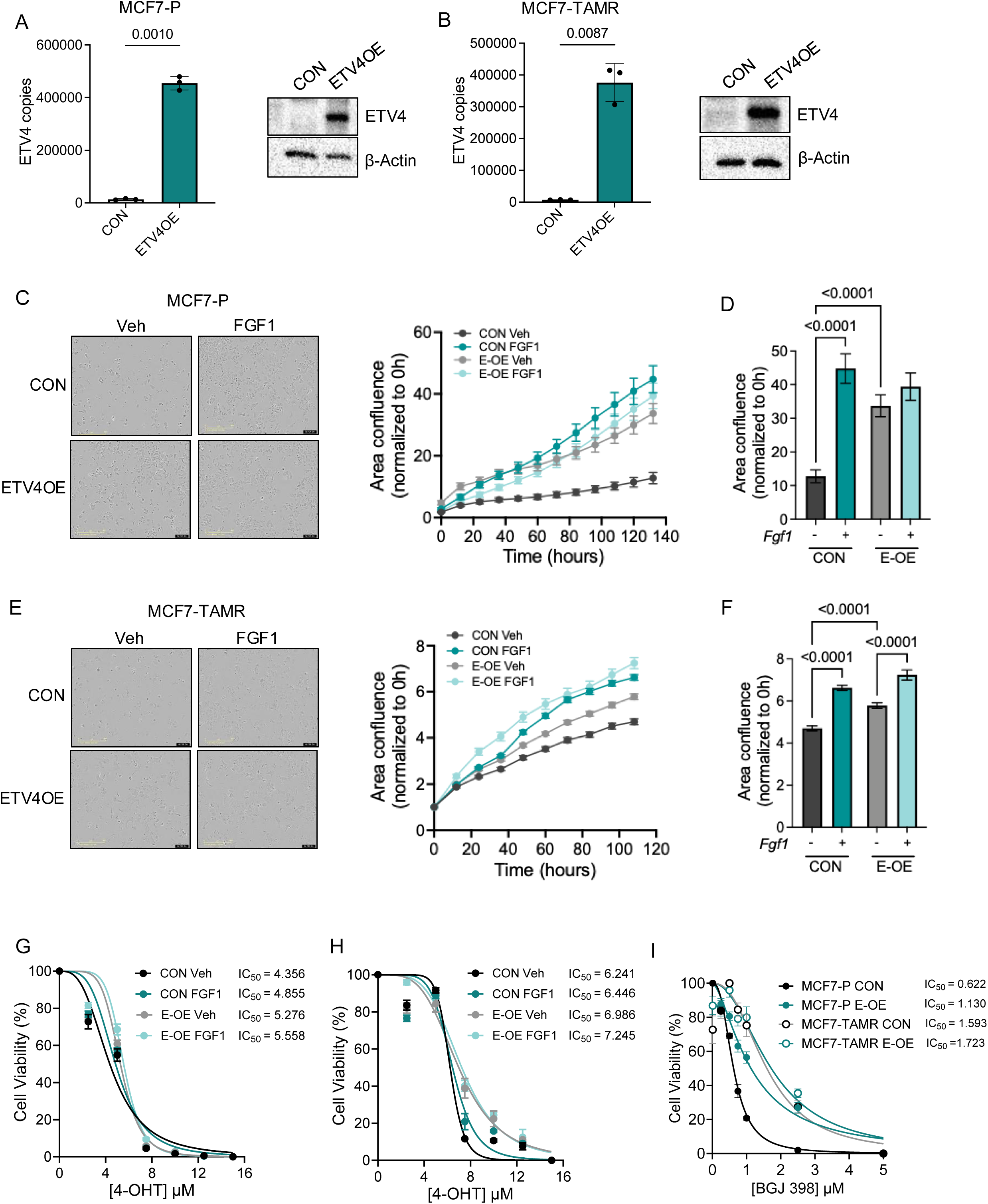
ETV4 overexpression promotes FGF1-mediated proliferation in ER-positive breast cancer cells. (A) Expression of ETV4 gene (*left*) and protein (*right*) following knockdown in MCF7-P cells. (B) Expression of ETV4 gene (*left*) and protein (*right*) following knockdown in MCF7-TAMR cells. (C) Representative images of the final timepoint (left) and growth rates (right) of MCF7-P control or ETV4-overexpressing cells treated with vehicle or FGF1. (D) Area confluence relative to control vehicle time 0 of cells at the final timepoint following treatment of MCF7-P cells. Two-way ANOVA testing for main effects of ETV4 overexpression or FGF1 treatment or interaction was performed. P-values indicate post-hoc multiple testing for specific differences between pre-defined comparisons. (E) Representative images of the final timepoint (left) and growth rates (right) of MCF7-TAMR control or ETV4-overexpressing cells treated with vehicle or FGF1. (F) Area confluence relative to control vehicle time 0 of cells at the final timepoint following treatment of MCF7-TAMR cells. Two-way ANOVA testing for main effects of ETV4 overexpression or FGF1 treatment or interaction was performed. P-values indicate post-hoc multiple testing for specific differences between pre-defined comparisons. (G) Dose-response curve and interpolated IC_50_ values of tamoxifen treatment in MCF7-P control and ETV4 overexpressing cells following treatment with or without FGF1. (H) Dose-response curve and interpolated IC_50_ values of tamoxifen treatment in MCF7-TAMR control and ETV4 overexpressing cells following treatment with or without FGF1. (I) Dose-response curve and interpolated IC_50_ values of BGJ398 treatment in MCF7-P and MCF7-TAMR control and ETV4 overexpressing cells.

### ETV4 and FGF1 influence cancer hallmarks

To expand our analysis of ETV4 effects in breast cancer, we performed global gene expression profiling using microarray in both ETV4 knockdown and overexpressing MCF7 TAMR cells with or without FGF1 stimulation. Gene set enrichment analysis (GSEA) of the whole transcriptome using the Molecular Signatures Database confirmed that knockdown of ETV4 dysregulated critical pathways in breast cancer cells, such as glycolysis, hypoxia, and epithelial to mesenchymal transition (EMT), even in the presence of FGF1 (Fig 4A; Additional File 2). Conversely, ETV4 overexpression in these cells enriched gene sets related to glycolysis, hypoxia, inflammatory response, and EMT (Fig 4A). Next, we evaluated the differentially expressed genes in ETV4-knockdown and overexpressing genes using the established cancer hallmarks gene sets [34]. ETV4-knockdown cells had significantly lower levels of genes associated with Reprogramming Energy Metabolism and Sustained Angiogenesis (Fig 4B; Additional File 3). Genes that increased in ETV-knockdown cells (i.e. genes that are repressed by ETV4) represented the Genomic Instability hallmark (Supplemental Fig 3). No cancer hallmarks were significantly associated with ETV4 overexpression at baseline; however, addition of FGF1 to ETV4-overexpressing cells enriched the Sustaining Proliferative Signaling hallmark (Fig 4C; Additional File 3). Together, these data highlight a broad role for ETV4 in regulating transcriptional programs associated with ER-positive cancer progression, particularly in the context of FGF1 exposure.

**Figure 4.**
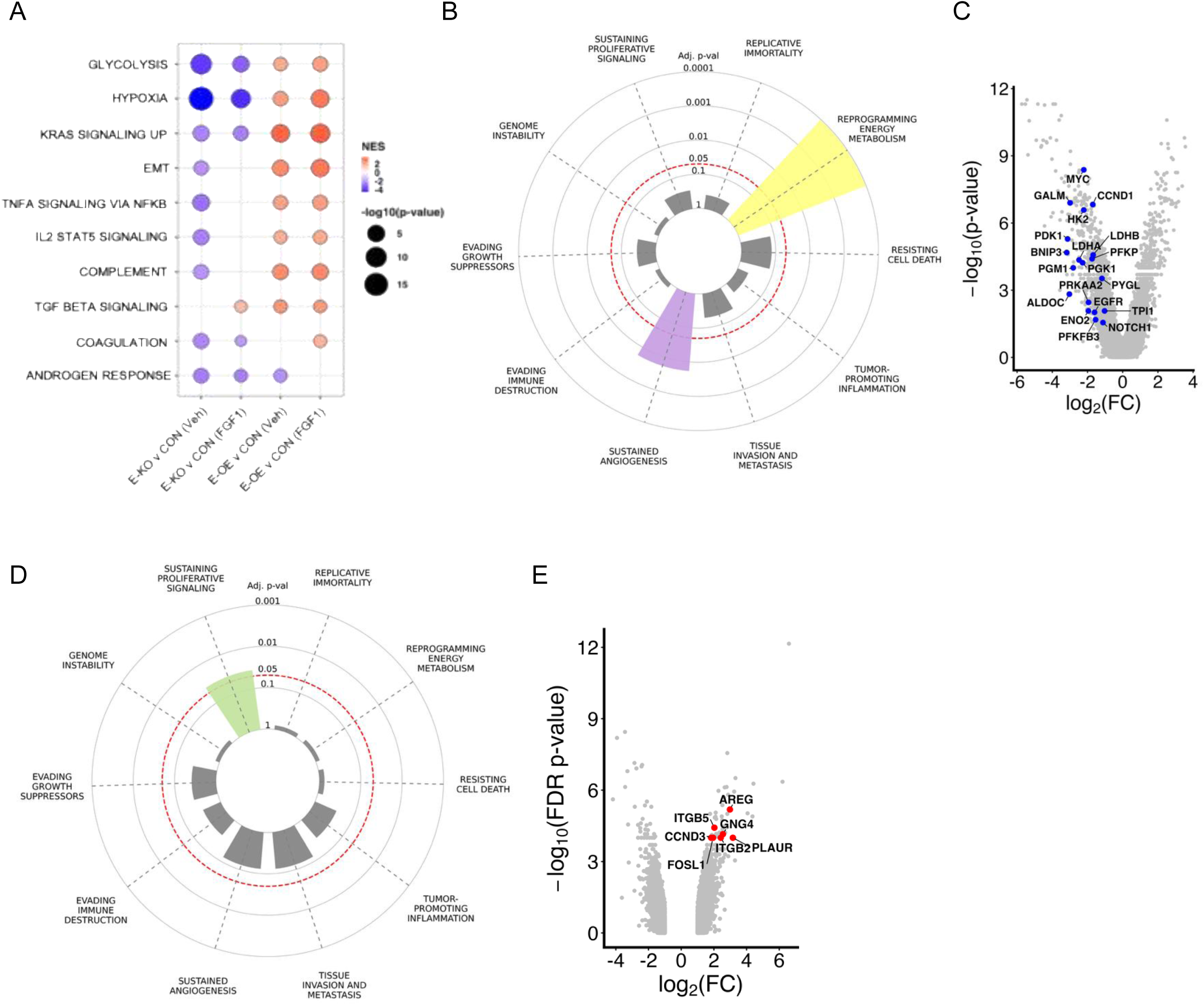
ETV4 and FGF1 alter genes associated with metabolism and aggressive breast cancer. (A) Bubble plot of gene set enrichment analysis (GSEA) showing enriched pathways in MCF7-TAMR control, ETV4 knockdown, and ETV4-overexpressing cells with or without FGF1 stimulation. (B) Hallmarks of Cancer enrichment plot illustrating the pathways represented by genes that are downregulated by ≥2-fold (adjusted p-value) in ETV4 knockdown compared with control vehicle-treated MCF7-TAMR cells. Bar height reflects −log10 adjusted p-value, with dashed circles indicating significance thresholds. (C) Volcano plot comparing ETV4 knockdown versus control vehicle-treated MCF7-TAMR cells, highlighting downregulated genes involved in reprogramming energy metabolism. Blue points denote significantly downregulated genes. Blue points denote significantly downregulated genes that correspond to the significant cancer hallmarks. (D) Hallmarks of Cancer enrichment plot illustrating the pathways represented by genes that are upregulated by ≥1.58-fold (adjusted p-value) in MCF7-TAMR ETV4 overexpressing cells treated with FGF1 vs vehicle controls. Bar height reflects −log10 adjusted p-value, with dashed circles indicating significance thresholds, including sustaining proliferative signaling. (E) Volcano plot comparing genes significantly altered in MCF7-TAMR ETV4 overexpressing cells treated with or without FGF1. Red points denote significantly upregulated genes that correspond to the significant cancer hallmarks.

### ETV4 regulates glycolysis in response to FGF1

FGF1 treatment promotes a glycolytic phenotype in ER-positive cells, measured by extracellular acidification rate (ECAR) and elevated glycolytic gene expression [16], therefore we measured these outcomes in control and ETV4 knockdown MCF7-P, MCF7-TAMR, and UCD12 models. In control MCF7-P cells, FGF1 induced rate-limiting genes in the glycolytic pathway, including HK2, PFKP, PGK1, ENO1, and LDHA (Fig 5A; Additional File 4). Knockdown of ETV4 had no effect on levels of these genes in the absence of FGF1; however, FGF1 treatment of ETV4 knockdown cells significantly increased expression of each gene (Fig 5A; Additional File 4). Treatment of control MCF7-TAMR (Fig 5B) and UCD12 cells (Supplemental Fig 4A) with FGF1 similarly induced expression of each gene, but in contrast to MCF7-P cells, the magnitude of induction by FGF1 in ETV4 knockdown cells was blunted. These results show that, although FGF1 induces ETV4 and glycolytic genes in multiple breast cancer cells, the signaling network connecting FGF1 to transcriptional regulation of these genes through ETV4 is not conserved across all ER-positive breast cancer models.

**Figure 5.**
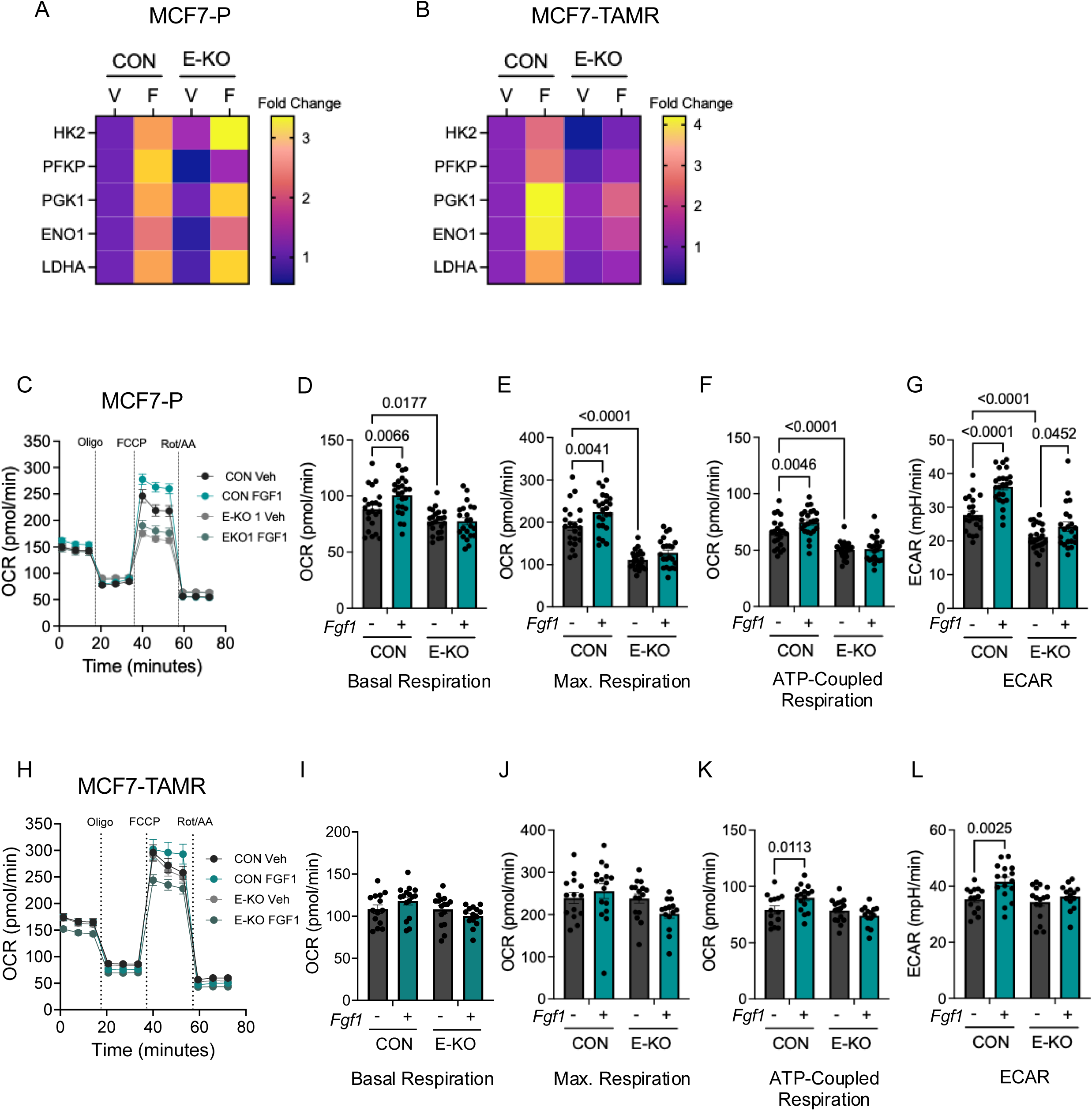
Loss of ETV4 suppresses FGF1-mediated glycolysis in breast cancer cells. (A-B) Heatmaps showing mRNA expression levels of glycolytic pathway genes (HK2, PFKP, PGK1, ENO1, and LDHA) under vehicle and FGF1-treated conditions in MCF7-P (*a*) and MCF7-TAMR (*b*) control and ETV4 knockdown cells, respectively. Data are expressed as fold change versus the average of vehicle treated cells for each gene, showing 3 replicates per group. (C) Seahorse metabolic flux analysis showing the kinetic graph of oxygen consumption rate (OCR) in MCF7-P cells. (D-G) Metabolic parameters including basal respiration (*d*), maximal respiration (*e*), ATP-production coupled respiration (*f*), and ECAR (*g*) in control and ETV4 knockdown MCF7-P cells, upon vehicle and FGF1 stimulation. Data analyzed using a 2-way ANOVA testing for main effects of ETV4 or FGF1 treatment or interactions. P-values denote post-hoc analysis of specific comparisons. (H) Seahorse metabolic flux analysis showing the kinetic graph of oxygen consumption rate (OCR) in MCF7-TAMR cells. (I-L) Metabolic parameters including basal respiration (*i*), maximal respiration (*j*), ATP-production coupled respiration (*k*), and ECAR (*l*) in control and ETV4 knockdown MCF7-TAMR cells, upon vehicle and FGF1 stimulation. Data were analyzed using a 2-way ANOVA testing for main effects of ETV4 or FGF1 treatment or interactions. P-values denote post-hoc analysis of specific comparisons. All Seahorse data were normalized to total protein in each well. N=16-24 replicates per measure.

Next, we investigated metabolic endpoints in cells with and without FGF1 treatment or ETV4, using the surrogate measures of oxygen consumption (OCR) and extracellular acidification rates (ECAR). In MCF7-P control cells, treatment with FGF1 and modulation of ETV4 expression impacted cellular oxygen consumption (Fig 5C). Basal and ATP-coupled OCR, which are calculated by subtracting non-mitochondrial oxygen consumption, as well as ECAR were significantly higher with FGF1 treatment (Fig 5D-G). Under baseline conditions (i.e. without FGF1 treatment), ETV4 knockdown cells had lower ATP-coupled respiration and ECAR compared with controls, and treatment with FGF1 did not increase any metabolic measure (Fig 5D-G). Oxygen consumption in MCF7-TAMR cells was also affected by FGF1 and ETV4 (Fig 5H), but to a lesser extent than MCF7-P cells. FGF1 treatment did not affect basal or maximal OCR (Fig 5I-J), but ATP-coupled OCR and ECAR increased with FGF1 in control cells (Fig 5K-L). In contrast to MCF7-P cells, the MCF7-TAMR cells were unaffected by ETV4 knockdown in the absence of FGF1 (Fig 5I-L). Like MCF7-P cells, FGF1 was ineffective at increasing any metabolic measure in the absence of ETV4 (Fig 5I-L). We also evaluated how ETV4 integrated the effects of FGF1 in UCD12 cells. Knocking down ETV4 lowered basal, maximal, and ATP-coupled respiration, but did not impact ECAR (Supplemental Fig 4B-F). Like both MCF7 lines, FGF1 increased ECAR in UCD12 controls, but did not increase metabolic activity of ETV4 knockdown cells (Supplemental Fig 4F). Together, these data reveal differences between glycolytic gene expression and functional outcomes across ER-positive breast cancer cell lines. While ETV4 is required for FGF1-dependent upregulation of glycolytic gene expression in two of the three models, the dependence of metabolic activity on ETV4 downstream of FGF1 occurs in each cell line, demonstrating that gene expression is not directly related to metabolic function.

### ETV4 is sufficient to increase glycolytic activity in ER-positive breast cancer cells

Based on the upregulation of ETV4 by FGF1 across breast cancer models, we predicted that ETV4 overexpression would be sufficient to promote metabolic reprogramming *in vitro*, as we have seen with FGF1 treatment. Consistent with the ETV4 knockdown experiments, FGF1 induced the expression of glycolytic genes in control MCF7-P cells (Fig 6A; Additional File 4). ETV4 overexpression alone also increased levels of these genes, and treatment with FGF1 further increased HK2, PFKP, and PGK1 expression in both cell lines (Fig 6A-B). ETV4 overexpression impacted metabolic activity at baseline and in the presence of FGF1 (Fig 6C). In control MCF7-P cells, FGF1 increased basal, maximal, and ATP-coupled respiration as well as ECAR (Fig 6D-G). ETV4 overexpression similarly increased each of these endpoints (Fig 6D-G). Basal (Fig 6D) and ATP-coupled respiration (Fig 6F), as well as ECAR (Fig 6G) were further elevated with FGF1 treatment in ETV4-overexpressing MCF7-P cells. In MCF7-TAMR cells, ETV4 overexpression was sufficient to increase glycolytic gene expression, which was further induced by FGF1 treatment (Fig 6G; Additional File 4). The effects of ETV4 overexpression on metabolic activity were similar to those seen in MCF7-P cells (Fig 6H). In MCF7-TAMR cells, ETV4 overexpression induced all metabolic measures (Fig 6I-L). Interestingly, FGF1 treatment only potentiated ECAR in cells with excess ETV4 (Fig 6L), without affecting oxygen consumption at any point. Together, these data reinforce the conclusion that induction of glycolytic genes is not directly tied to metabolic activity in ER-positive breast cancer cells, but that FGF1 influences functional changes in metabolism, particularly extracellular acidification, in cooperation with ETV4.

**Figure 6.**
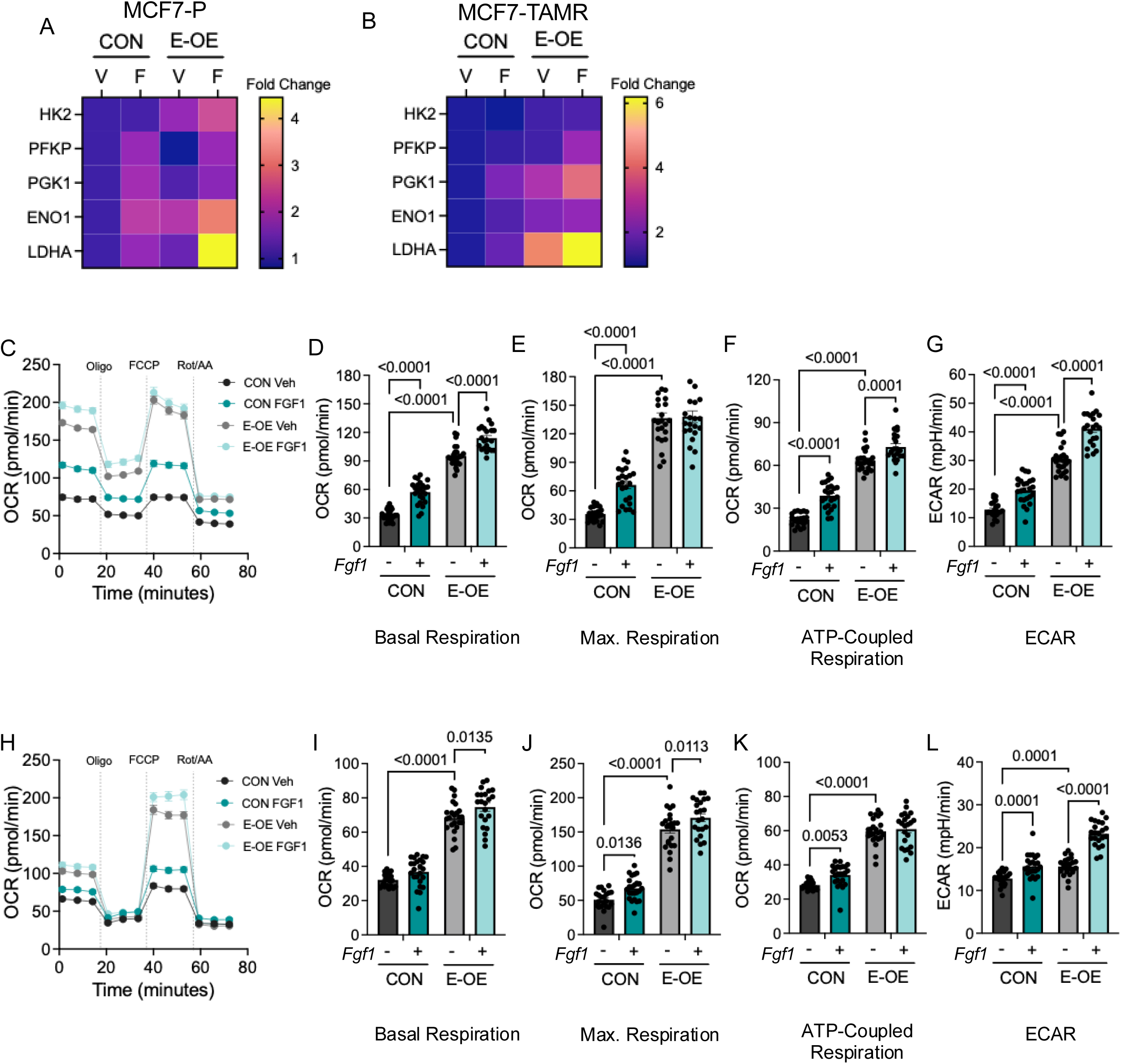
ETV4 overexpression enhances FGF1-mediated glycolysis in breast cancer cells. (A-B) Heatmaps showing mRNA expression levels of glycolytic pathway genes (HK2, PFKP, PGK1, ENO1, and LDHA) under vehicle and FGF1-treated conditions in MCF7-P (*a*) and MCF7-TAMR (*b*) control and ETV4 overexpressing cells, respectively. Data are expressed as fold change versus the average of vehicle treated cells for each gene, showing 3 replicates per group. (C) Seahorse metabolic flux analysis showing the kinetic graph of oxygen consumption rate (OCR) in MCF7-P cells. (D-G) Metabolic parameters including basal respiration (*d*), maximal respiration (*e*), ATP-production coupled respiration (*f*), and ECAR (*g*) in control and ETV4 overexpressing MCF7-P cells upon vehicle and FGF1 stimulation. Data were analyzed using a 2-way ANOVA testing for main effects of ETV4 or FGF1 treatment or interactions. P-values denote post-hoc analysis of specific comparisons. (H) Seahorse metabolic flux analysis showing the kinetic graph of oxygen consumption rate (OCR) in MCF7-TAMR cells. (I-L) Metabolic parameters including basal respiration (*i*), maximal respiration (*j*), ATP-production coupled respiration (*k*), and ECAR (*l*) in control and ETV4 knockdown MCF7-TAMR cells, upon vehicle and FGF1 stimulation. Data were analyzed using a 2-way ANOVA testing for main effects of ETV4 or FGF1 treatment or interactions. P-values denote post-hoc analysis of specific comparisons. All Seahorse data were normalized to total protein in each well. N=16-24 replicates per measure.

### ETV4 is necessary but not sufficient for ER-positive tumor growth

Our previous studies have shown that obesity supports estrogen-independent growth of some, but not all ER-positive tumors [13, 16]. Specifically, we reported that MCF7-TAMR and UCD12 tumors continued to progress in obese mice after estrogen withdrawal (EWD), but MCF7-P tumors did not. The data presented here reveal consistent regulation of ETV4 expression, cell proliferation, and metabolic activity downstream of FGF1 in breast cancer cell lines, but effects *in vitro* do not always translate to relevant impacts on tumor growth *in vivo*. We reasoned that, if obesity-associated FGF1 upregulates ETV4 to support tumor progression, then ETV4 overexpression would be sufficient to permit tumor growth after EWD in lean mice, where FGF1 levels are not elevated. To test this, we grafted control or ETV4 overexpressing MCF7-TAMR cells into inguinal mammary fat pads of lean females (Fig 7A). Estradiol (E2) was administered at the time of tumor implantation, and tumor growth was measured. Once tumors reached a specified volume (∼800 mm^3^), E2 was withdrawn to mimic endocrine therapy, and tumor endpoints were measured after 4 weeks (Fig 7A).

**Figure 7.**
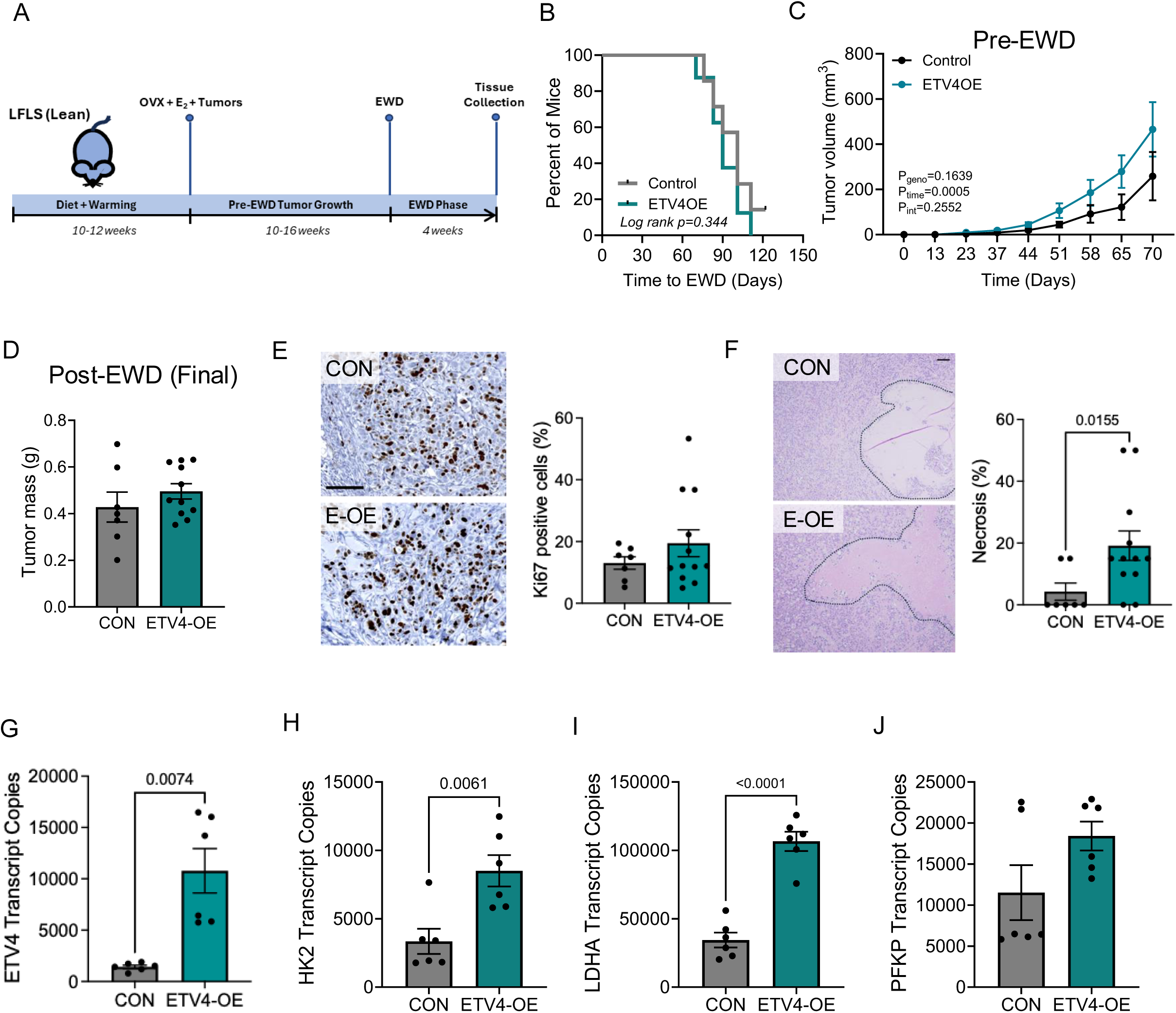
ETV4 overexpression is not sufficient to support ER-positive tumor growth in lean mice. (A) Schematic diagram showing the experimental timeline. Female mice were maintained under low-fat, low-sucrose (LFLS; lean) diet conditions for 10-12 weeks, followed by ovariectomy (OVX) and supplementation with estradiol (E₂) at the time of tumor implantation. Tumors grew for 10-16 weeks prior to estrogen withdrawal (EWD), which occurred when the largest tumor per mouse reached ∼800mm^3^. EWD was performed for 4 weeks, after which tumors and tissues were collected for downstream analyses. (B) Kaplan-Meier survival analysis showing time to EWD for mice grafted with MCF7-TAMR control or ETV4 overexpressing cells. (C) Tumor volumes in mice bearing control or ETV4-overexpressing cells. Growth is plotted from the time of tumor implantation until the first mouse in either group was assigned to EWD treatment. Data were analyzed using 2-way ANOVA, testing for main effects of time or ETV4 expression, or interactions. (D) Tumor mass in mice after 4 weeks of EWD treatment. (E) Representative IHC images of Ki67 staining from tumor samples post-EWD and corresponding quantification of Ki67-positive cells. Scale = 100 µm. (F) Representative images of H&E-stained tumors following EWD, with quantification of necrosis represented as a percentage. Outlines denote necrotic regions. Scale = 50 µm. (G-J) Expression of ETV4 (*g*) and glycolytic genes HK2 (*h*), LDHA (*i*), and PFKP (*j*) in tumor samples after EWD.

In the presence of supplemental E2, which is necessary to establish human ER-positive tumors in mice, ETV4 overexpression had no impact on tumor parameters. Compared with control tumors, there were more injected tumors that successfully grafted in the ETV4-overexpressing group (i.e. higher “take rate”), but the differences were not significant (Supplemental Fig 5A). There was no effect of ETV4 on the time to reach EWD (Fig 7B) or on the growth rates of individual tumors prior to EWD (Fig 7C; Supplemental Fig 5B-C). To determine whether ETV4 could sustain tumors in the lean environment, we withdrew supplemental E2 once they were established. Irrespective of ETV4 overexpression, tumors regressed after EWD and there was no difference in final tumor mass or volume between groups (Fig 7D; Supplemental Fig 5D). ETV4-overexpressing tumors had a slightly, but non-significantly higher Ki67 index compared to controls at the end of the study (Fig 7E), and they had significantly more necrosis (Fig 7F). Further analysis revealed that the ETV4 overexpressing tumors maintained high ETV4 levels (Fig 7G) and had elevated expression of HK2 (Fig 7H) and LDHA (Fig 7I), but there was no significant difference in PFKP levels (Fig 7J). These data suggest that, although ETV4 overexpression *in vitro* is sufficient to drive proliferation, glycolytic gene expression, and greater metabolic activity, it is not sufficient to accelerate tumor growth before or after EWD in lean mice.

Following the same rationale, we evaluated whether ETV4 was necessary to support tumor growth in the obese environment (Fig 8A), where we have documented elevated FGF1 production [13]. ETV4 knockdown reduced the number of grafted tumors that successfully established (Supplemental Fig 6A), although this was not significant. Tumors with reduced ETV4 were slower to reach the EWD treatment volume (Fig 8B) and were significantly smaller at the time when control mice underwent EWD treatment (Fig 8C). However, once ETV4 knockdown tumors formed, their growth rates were similar to controls (Supplemental Fig 6B-C). After EWD, there were no significant differences in tumor mass (Fig 8D) or volume (Supplemental Fig 6D) between groups. Consistent with this, the Ki67 index and necrosis regions were comparable between control and ETV4-knockdown tumors (Fig 8E-F). ETV4 reduction was confirmed in tumors at the end of the study (Fig 8G). HK2 was unaffected by ETV4 knockdown, but PFKP and LDHA expression were lower (Fig 8H-J). Together, these studies indicate that ETV4 is not sufficient to promote ER-positive tumor growth in lean females regardless of E2 supplementation. In obese females, ETV4 partially mediates E2-dependent tumor growth; however, once estrogen is depleted, the impact of ETV4 on tumor progression is minimal.

**Figure 8.**
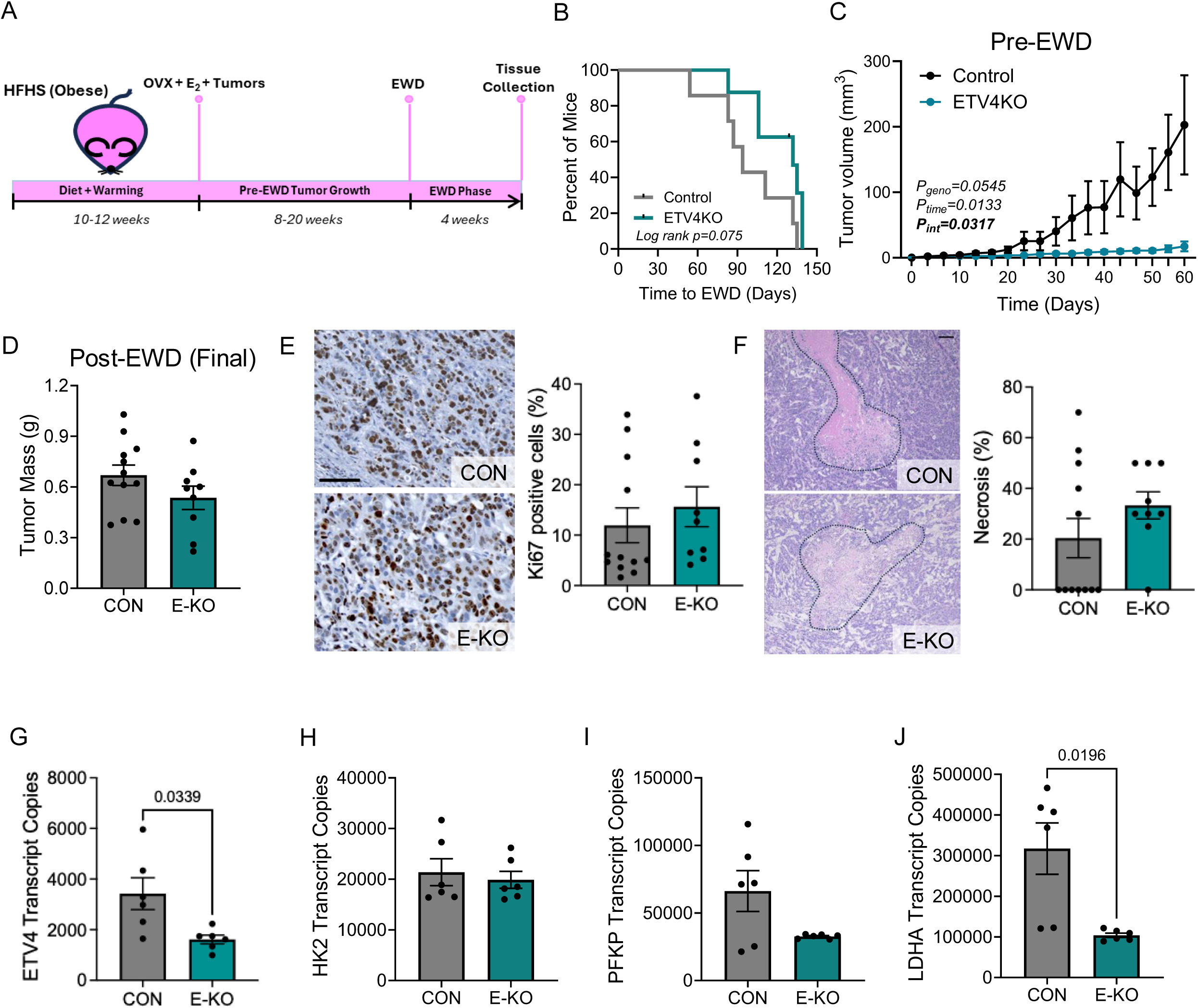
ETV4 loss delays tumor growth in obese mice. (A) Schematic diagram showing the experimental timeline. Female mice were maintained under high-fat, high-sucrose (HFHS; obese) diet conditions for 10-12 weeks, followed by ovariectomy (OVX) and supplementation with estradiol (E₂) at the time of tumor implantation. Tumors grew for 8-20 weeks prior to estrogen withdrawal (EWD). EWD was performed for 4 weeks, after which tumors and tissues were collected for downstream analyses. (B) Kaplan-Meier survival analysis showing time to EWD for mice grafted with MCF7-TAMR control or ETV4 knockdown cells. (C) Tumor volumes in mice bearing control or ETV4-knockdown cells. Growth is plotted from the time of tumor implantation until the first mouse in either group was assigned to EWD treatment. Data were analyzed using 2-way ANOVA, testing for main effects of time or ETV4 expression, or interactions. (D) Tumor mass in mice after 4 weeks of EWD treatment. (E) Representative IHC images of Ki67 staining from tumor samples post-EWD and corresponding quantification of Ki67-positive cells. Scale = 100 µm. (F) Representative images of H&E-stained tumors following EWD, with quantification of necrosis represented as a percentage. Outlines denote necrotic regions. Scale = 50 µm. (G-J) Expression of ETV4 (*g*) and glycolytic genes HK2 (*h*), PFKP (*i*), and LDHA (*j*) in tumor samples after EWD.

## DISCUSSION

We carried out this study to extend our work on obesity and ER-positive breast cancer progression. Previously, we identified FGF1 as a potential driver of estrogen-independent tumor growth in obese female rats and mice, and reported that blocking FGFR signaling restored sensitivity of ER-positive tumors to estrogen deprivation *in vivo* [13, 14]. We also demonstrated that, although FGF1 treatment had pro-tumorigenic effects on gene expression and metabolism in ER-positive breast cancer cells, not all cell lines were sustained by obesity (i.e. excess FGF1) when grown as orthotopic tumors [16]. These observations are expected given that some patients with obesity do not have an elevated risk for ER-positive breast cancer diagnosis or progression, highlighting the heterogeneity within this tumor subtype. ETV4 emerged as one of the top genes consistently induced by FGF1 *in vitro*, thus our goal was to determine to what extent ETV4 was necessary or sufficient to mediate effects of, or substitute for FGF1 exposure in breast cancer cells.

Here, we show that ETV4 expression is higher in tumors compared with normal breast tissue and is also associated with shorter overall and disease-free survival of people with ER-positive breast cancer. Elevated ETV4 levels have been reported in mouse, rat, and human breast tumors [19, 35, 36], and predict a poor prognosis [23, 28]. We now extend these findings to demonstrate that high ETV4 expression is more prevalent in ER-negative tumors, which have a relatively poor prognosis and can complicate analyses of cancer mortality when combined with ER-positive breast cancer. When we focused only on ER-positive tumors, high ETV4 levels remained a negative prognostic indicator. While expression of ETV4 alone was not associated with BMI in primary human breast cancer specimens, the positive correlation between ETV4 and FGF1 was seen only in tumors from people with obesity. FGF1 upregulated ETV4 expression in multiple ER-positive breast cancer cell lines, which is aligned with data from others showing that FGF2 induced ETV4 in MDA-231 cells and promoted ETV4 binding to Ets-response elements in MCF7 cells [28]. Importantly, we previously reported that FGF1, but not FGF2 was coordinately upregulated in mammary adipose during estrogen withdrawal-induced weight gain in obese mice [13]. By focusing the studies here on FGF1, we reveal cellular mechanisms connecting weight gain and obesity to breast cancer progression. ETV4 with and without FGF1 also influenced the sensitivity of cells to tamoxifen, which is a standard treatment for breast cancer. Tamoxifen treatment was shown to reduce ETV4 expression in normal breast samples from women at high risk for breast cancer [37], suggesting the potential for further development of ETV4 as a predictive biomarker of pro- and anti-tumorigenic changes in the breast.

Transcriptomics analysis revealed a significant downregulation of several oncogenic pathways, including hypoxia, epithelial to mesenchymal transition (EMT), glycolysis, and inflammatory response, upon ETV4 loss. The cancer hallmark, Reprogramming Energy Metabolism, was also markedly impacted by the loss of ETV4. Conversely, ETV4 overexpression enriched similar pathways, especially in the presence of FGF1. Others have shown that ETV4 regulates genes involved in cell cycle regulation, EMT, invasion, and metastasis in cancer cells [38–42], and data presented here demonstrate that this also occurs in models of ER-positive breast cancer. ETV4 is reported to regulate glycolytic gene expression and activity, supporting the maintenance of stem cell pools in triple-negative breast cancer cells [21]. While we did not evaluate breast cancer stem cells, our validation of key glycolytic genes in cells with and without ETV4 revealed interesting differences across models. In MCF7-P cells, FGF1 induced several glycolytic enzymes regardless of ETV4 presence; however, in MCF7-TAMR and UCD12 cells, the effect of FGF1 was attenuated upon ETV4 loss. In all three ER-positive breast cancer cell lines, we showed that FGF1 increases glycolysis, inferred from ECAR [16], and here we show that this is mediated by ETV4. Reduction of ETV4 lowered cellular oxygen consumption, and the FGF1-dependent increase in glycolysis was consistently attenuated. Conversely, ETV4 overexpression increased oxidative metabolism, and further elevated glycolytic activity in the presence of FGF1. Therefore, despite differences in the dependence of FGF1-induced glycolytic gene expression on ETV4 between cell lines, the functional experiments repeatedly demonstrated a role for ETV4 in mediating elevated metabolic activity on its own and downstream of FGF1. These data suggest that the network connecting FGF1 and ETV4 transcriptional activity to glycolytic gene expression is cell-line dependent, but the transcriptomic changes do not always correlate with biological function.

To extend the *in vitro* studies and better understand how ETV4 influences tumor progression, we performed *in vivo* experiments in our established preclinical model [13, 16, 43]. We hypothesized that obesity promotes estrogen-independent cancer progression through FGF1-mediated induction of ETV4. Therefore, ETV4 overexpression should be sufficient to permit tumor growth after EWD in lean mice, where FGF1 levels remain low [13]. Conversely, tumors with reduced ETV4 expression should not grow after estrogen deprivation in obese mice, even though FGF1 should increase during EWD-induced weight gain. We chose MCF7-TAMR cells for these experiments because MCF7-P cells are critically dependent upon supplemental estrogen and do not grow in mice after EWD, regardless of host adiposity [13]. The most striking effects were seen in the presence of estradiol (E2). ETV4 overexpressing tumors were indistinguishable from controls in lean mice, whereas growth of ETV4 knockdown tumors was attenuated over time in obese mice; however, this only occurred prior to EWD. Contrary to what we expected based on *in vitro* data, after EWD, ETV4 overexpression was not sufficient to sustain tumors in lean females, nor was it required to maintain tumors in obesity. Thus, despite the consistent and substantial role for ETV4 in cultured breast cancer cell proliferation, gene expression, and metabolic activity, additional variables might underlie the progression of ER-positive tumors *in vivo* and in humans, where ETV4 levels did associate with response to therapy.

In summary, our study provides new insights into the potential for targeting the FGF1/ETV4 axis in obesity-associated ER-positive breast cancer. We suggest that ETV4 is a critical transcription factor that acts downstream of FGF1 signaling to regulate cell proliferation and rewires metabolism to drive aggressive breast cancer phenotypes, which may be more prevalent in the context of obesity. Future studies should involve greater emphasis on factors beyond BMI that influence the response of ER-positive breast cancer to therapy and the patients’ long-term outcomes.

## MATERIALS & METHODS

### Cell lines and reagents

MCF7 and tamoxifen-resistant MCF7 (TAMR) cells were purchased from America Type Cell Collection (ATCC, USA) and were maintained in Dulbecco’s modified Eagle’s medium (DMEM) supplemented with 10% fetal bovine serum (FBS), 1% penicillin/streptomycin (P/S), and 10 μg/mL insulin at 37°C in the presence of 5% CO₂. UCD12 cells were cultured in DME/F12 with 10% FBS, 1% pen/strep, 100 ng/mL cholera toxin, and 1 nM insulin. During experimental treatments, cells were starved for 16 hours in phenol red-free Minimum Essential Medium (MEM) containing 0.5% charcoal-stripped fetal bovine serum to decrease the potential of steroid hormone receptor activation. All growth factor treatments were given in this medium. Recombinant human FGF1 was purchased from R&D Systems and was diluted in 0.01% bovine serum albumin (BSA) in PBS and used at a final concentration of 5 ng/mL. All cell lines were routinely tested for mycoplasma using the MycoAlert Mycoplasma Detection kit (Lonza, Rockland, ME, USA).

### Stable cell line generation

To generate ETV4 stable knockdown cell lines, lentiviral particles equipped with short hairpin RNAs (shRNAs) targeting ETV4 were purchased from Santa Cruz (sc-36205-V) and Millipore Sigma (clone TRCN0000013937) as was a non-specific shRNA control (sc-108080). Following the manufacturer’s instructions, cells were transduced with viral particles and screened for stable clones using 2 μg/ml puromycin. ETV4-overexpressing cells were generated using lentiviral plasmid pLX_TRC317 (Addgene #142031), and stable clones were selected using puromycin. The expression of ETV4 was determined by western blot and quantitative RT-PCR.

### Cell proliferation assay

The InCucyte S3-Live Cell Analysis instrument was used to perform cell proliferation. Briefly, 2 x 10^3^ cells were seeded in a 96-well plate with or without FGF1 treatment and monitored as indicated in each figure. Live cell images were obtained at a 10x objective lens, and area confluence was estimated.

### Cytotoxicity Assay

Cells were seeded in 96-well plates in triplicate in complete growth media. The next day, the media was changed to hormone-free media containing varying concentrations of (0-20 μM) 4-Hydroxytamoxifen or (0-10 μM) FGFR1 inhibitor BGJ398 in the presence or absence of FGF1 for 72 hours. Cytotoxicity was assessed using the CellTitre Glo 2D Cell Viability Assay (Promega) on day 3 according to the manufacturer’s instructions. GraphPad Prism 10.4 software was used to determine the half-maximal inhibitory concentration (IC_50_) value for each cell line.

### Metabolic flux analysis

The Seahorse Mito Stress test was used to measure basal, ATP-linked, and maximal oxygen consumption (OCR) and extracellular acidification rate (ECAR) according to the manufacturer’s instructions (Cat. No. 103015-100). Briefly, cells were seeded into Seahorse XF96 cell culture microplates at a density of 7000 cells/well. After 24 hours, cells were treated with either 5 ng/ml FGF1 or EtOH (vehicle) overnight. On the day of assay, cells were incubated in XF DMEM medium supplemented with 1 mM D-glucose, 2 nM L-glutamine, and 1 mM sodium pyruvate for 1 hour before the test in a non-CO₂ incubator at 37°C. Next, the cells were stimulated with 1.5 μM oligomycin, 1 μM FCCP, and 0.5 μM Rotenone/Antimycin A to measure oxygen consumption and extracellular acidification rate. The data were normalized to cell number using the Sulforhodamine B colorimetric assay.

### Human dataset analysis

The TNM plot database was used to analyze ETV4 expression in normal breast tissue and invasive breast tumors [30]. The KM plotter database was used to analyze the association between ETV4 mRNA expression and overall or recurrence-free survival in breast tumors, regardless of subtype [44]. The ROC analysis database was used to evaluate the prognostic significance of ETV4 expression in ER-positive breast cancer, as well as the gene levels in tumors classified as responders or non-responders to therapy [31]. Correlation analysis was performed using raw data obtained from the microarray datasets GSE24185 [45] and GSE186901 [46] from the Gene Expression Omnibus database.

### Immunoblot analysis

For Western blot analysis, cells were lysed in RIPA buffer with protease inhibitors (Roche) and PhosSTOP phosphatase inhibitors (Roche). Protein concentration was measured by the Pierce bicinchoninic acid (BCA) Protein Assay kit (Thermo Fisher Scientific, Rockford, IL). Approximately 30-40 μg of protein from each sample was separated on an 8-16% Mini-PROTEAN TGX precast gel (Bio-Rad Lab Inc., USA) and transferred to a polyvinylidene fluoride membrane. The following primary antibodies were used: ETV4 (Cat. No. 10684-1-AP, Proteintech), Vinculin (Cat. No. 13901, Cell Signaling Technology), or β-Actin (Cat. No. 4970, Cell Signaling Technology), followed by HRP-linked anti-rabbit secondary antibodies (Cell Signaling Technology). Images were taken using the iBright™ FL1500 Imaging system and quantified with the ImageJ software (US National Institute of Health).

### Immunohistochemistry (IHC) staining

Formalin-fixed and paraffin-embedded breast tumor sections were stained with Hematoxylin and Eosin (H&E) and IHC was performed, using the standardized protocol at the Stephenson Cancer Center Tissue Pathology core. For IHC, the Ki-67 antibody (Cat. No. 122025, Cell Signaling Technology) was used at a dilution of 1:3000 for 60 minutes, followed by ImmPRESS® HRP Horse Anti Rabbit IgG Polymer Detection kit (Vector Laboratories, MP-7401). Stained slides were digitally scanned using the Zeiss Axio Scan at 20x magnification. Quantification was performed using the HALO software (Indica Lab, V3.2.1851). The IHC and H&E slides were independently scored by a pathologist.

### Quantitative PCR

Total RNA was extracted using the RNeasy Mini Kit (Qiagen, Hilden, Germany 74134). A total of 1 µg of total RNA was reverse transcribed into cDNA using the Verso cDNA synthesis kit (ThermoFisher, MA, USA). cDNA representing 25 ng of total RNA was added to each qPCR reaction containing TaqMan Fast Advanced Master Mix primers (ThermoFisher, MA, USA) specific to ETV4, HK2, PFKP, PGK1, ENO1, AND LDHA. Data are presented as transcript copy numbers, estimated using a standard curve of the amplicon as previously described [47].

### Microarray and bioinformatics

Total RNA from MCF7-TAMR control, ETV4 knockdown, and ETV4 overexpressing lines treated with or without FGF1 was submitted to North American Genomics, Inc., for global gene expression profiling using Human Clariom S Array. The data were analyzed with Transcriptome Analysis Console (TAC) 4.0.2 (Thermo Fisher Scientific) to identify the differentially expressed genes between the treatment groups. Multiple testing correction identified DEGs using an FDR cutoff of <0.05. The fgsea package in R was used to perform Gene Set Enrichment Analysis (GSEA) on the DEGs using the Hallmark gene set from the Molecular Signatures Database. The established Cancer Hallmarks were determined using cancerhallmarks.com [48]. Raw microarray data will be deposited in the NCBI Gene Expression Omnibus.

### Mouse studies

All animal procedures were approved by the IACUC at the University of Oklahoma Health Center. Female Rag1KO mice (stock #002216) were purchased from The Jackson Laboratories at 6 weeks old. Upon arrival, mice were placed on either a high-fat (40% kcal fat) or a low-fat (11% kcal fat) diet purchased from Research Diets, Inc., as previously described [13, 43]. At 18-20 weeks of age (10-12 weeks on diet), mice were ovariectomized and supplemented with 0.5 μM 17β-estradiol in drinking water [43]. MCF7-TAMR cells were orthotopically implanted in the inguinal mammary fat pads of low-fat-fed or high-fat-fed mice (1 x 10^6^ cells per gland). Tumor growth was monitored by weekly caliper measurements until the volume reached the pre-specified size for estrogen deprivation treatment. Once the largest tumor per mouse reached ∼800 mm³, supplemental estradiol was withdrawn, and studies were terminated 4 weeks later. Mice were fasted for 4 hours prior to euthanasia and tumors were collected.

### Statistical analyses

Results are presented as mean ± SEM, with p-values less than 0.05 considered significantly different. Student t-test, multiple comparison t-test, one-way ANOVA, two-way ANOVA, and Pearson correlation were performed as indicated using GraphPad Prism software version 10.6.1 and R V4.4.3.

## REFERENCES

1. Siegel RL, Kratzer TB, Giaquinto AN, Sung H, Jemal A: Cancer statistics, 2025. CA Cancer J Clin 2025, 75(1):10–45.

2. Kennecke H, Yerushalmi R, Woods R, Cheang MC, Voduc D, Speers CH, Nielsen TO, Gelmon K: Metastatic behavior of breast cancer subtypes. J Clin Oncol 2010, 28(20):3271–3277.

3. Burstein HJ, Somerfield MR, Barton DL, Dorris A, Fallowfield LJ, Jain D, Johnston SRD, Korde LA, Litton JK, Macrae ER et al: Endocrine Treatment and Targeted Therapy for Hormone Receptor-Positive, Human Epidermal Growth Factor Receptor 2-Negative Metastatic Breast Cancer: ASCO Guideline Update. J Clin Oncol 2021, 39(35):3959–3977.

4. García-Estévez L, Cortés J, Pérez S, Calvo I, Gallegos I, Moreno-Bueno G: Obesity and Breast Cancer: A Paradoxical and Controversial Relationship Influenced by Menopausal Status. Front Oncol 2021, 11:705911.

5. Lorincz AM, Sukumar S: Molecular links between obesity and breast cancer. Endocr Relat Cancer 2006, 13(2):279–292.

6. Picon-Ruiz M, Morata-Tarifa C, Valle-Goffin JJ, Friedman ER, Slingerland JM: Obesity and adverse breast cancer risk and outcome: Mechanistic insights and strategies for intervention. CA Cancer J Clin 2017, 67(5):378–397.

7. Suzuki R, Iwasaki M, Inoue M, Sasazuki S, Sawada N, Yamaji T, Shimazu T, Tsugane S: Body weight at age 20 years, subsequent weight change and breast cancer risk defined by estrogen and progesterone receptor status--the Japan public health center-based prospective study. Int J Cancer 2011, 129(5):1214–1224.

8. Bhardwaj P, Au CC, Benito-Martin A, Ladumor H, Oshchepkova S, Moges R, Brown KA: Estrogens and breast cancer: Mechanisms involved in obesity-related development, growth and progression. J Steroid Biochem Mol Biol 2019, 189:161–170.

9. Harborg S, Cronin-Fenton D, Jensen MR, Ahern TP, Ewertz M, Borgquist S: Obesity and Risk of Recurrence in Patients With Breast Cancer Treated With Aromatase Inhibitors. JAMA Netw Open 2023, 6(10):e2337780.

10. Sho M, Qureshi R, Slingerland J: Oestrogen changes at menopause: insights into obesity-associated breast risk and outcomes. Nat Rev Endocrinol 2025.

11. Ligibel JA, Strickler HD: Obesity and Its Impact on Breast Cancer: Tumor Incidence, Recurrence, Survival, and Possible Interventions. American Society of Clinical Oncology Educational Book 2013(33):52–59.

12. Elliott MJ, Ennis M, Pritchard KI, Townsley C, Warr D, Elser C, Amir E, Bedard PL, Rao L, Stambolic V et al: Association between BMI, vitamin D, and estrogen levels in postmenopausal women using adjuvant letrozole: a prospective study. NPJ Breast Cancer 2020, 6:22.

13. Wellberg EA, Kabos P, Gillen AE, Jacobsen BM, Brechbuhl HM, Johnson SJ, Rudolph MC, Edgerton SM, Thor AD, Anderson SM et al: FGFR1 underlies obesity-associated progression of estrogen receptor-positive breast cancer after estrogen deprivation. JCI Insight 2018, 3(14).

14. Wellberg EA, Corleto KA, Checkley LA, Jindal S, Johnson G, Higgins JA, Obeid S, Anderson SM, Thor AD, Schedin PJ et al: Preventing ovariectomy-induced weight gain decreases tumor burden in rodent models of obesity and postmenopausal breast cancer. Breast cancer research : BCR 2022, 24(1):42.

15. Wang S, Cao S, Arhatte M, Li D, Shi Y, Kurz S, Hu J, Wang L, Shao J, Atzberger A et al: Adipocyte Piezo1 mediates obesogenic adipogenesis through the FGF1/FGFR1 signaling pathway in mice. Nat Commun 2020, 11(1):2303.

16. Castillo-Castrejon M, Sankofi BM, Murguia SJ, Udeme AA, Cen HH, Xia YH, Thomas NS, Berry WL, Jones KL, Richard VR et al: FGF1 supports glycolytic metabolism through the estrogen receptor in endocrine-resistant and obesity-associated breast cancer. Breast cancer research : BCR 2023, 25(1):99.

17. Laudet V, Hänni C, Stéhelin D, Duterque-Coquillaud M: Molecular phylogeny of the ETS gene family. Oncogene 1999, 18(6):1351–1359.

18. Chotteau-Lelièvre A, Desbiens X, Pelczar H, Defossez P-A, de Launoit Y: Differential expression patterns of the PEA3 group transcription factors through murine embryonic development. Oncogene 1997, 15(8):937–952.

19. Kurpios NA, Sabolic NA, Shepherd TG, Fidalgo GM, Hassell JA: Function of PEA3 Ets transcription factors in mammary gland development and oncogenesis. J Mammary Gland Biol Neoplasia 2003, 8(2):177–190.

20. Yuan ZY, Dai T, Wang SS, Peng RJ, Li XH, Qin T, Song LB, Wang X: Overexpression of ETV4 protein in triple-negative breast cancer is associated with a higher risk of distant metastasis. Onco Targets Ther 2014, 7:1733–1742.

21. Zhu T, Zheng J, Zhuo W, Pan P, Li M, Zhang W, Zhou H, Gao Y, Li X, Liu Z: ETV4 promotes breast cancer cell stemness by activating glycolysis and CXCR4-mediated sonic Hedgehog signaling. Cell Death Discovery 2021, 7(1).

22. Shepherd TG, Kockeritz L, Szrajber MR, Muller WJ, Hassell JA: The pea3 subfamily ets genes are required for HER2/Neu-mediated mammary oncogenesis. Curr Biol 2001, 11(22):1739–1748.

23. Zhu T, Zheng J, Zhuo W, Pan P, Li M, Zhang W, Zhou H, Gao Y, Li X, Liu Z: ETV4 promotes breast cancer cell stemness by activating glycolysis and CXCR4-mediated sonic Hedgehog signaling. Cell Death Discov 2021, 7(1):126.

24. Garg A, Hannan A, Wang Q, Collins T, Teng S, Bansal M, Zhong J, Xu K, Zhang X: FGF-induced Pea3 transcription factors program the genetic landscape for cell fate determination. PLoS Genet 2018, 14(9):e1007660.

25. Garg A, Hannan A, Wang Q, Makrides N, Zhong J, Li H, Yoon S, Mao Y, Zhang X: Etv transcription factors functionally diverge from their upstream FGF signaling in lens development. Elife 2020, 9.

26. Zhang Z, Verheyden JM, Hassell JA, Sun X: FGF-regulated Etv genes are essential for repressing Shh expression in mouse limb buds. Dev Cell 2009, 16(4):607–613.

27. DeSalvo J, Ban Y, Li L, Sun X, Jiang Z, Kerr DA, Khanlari M, Boulina M, Capecchi MR, Partanen JM et al: ETV4 and ETV5 drive synovial sarcoma through cell cycle and DUX4 embryonic pathway control. The Journal of clinical investigation 2021, 131(13).

28. Myers E, Hill AD, Kelly G, McDermott EW, O’Higgins NJ, Young LS: A positive role for PEA3 in HER2-mediated breast tumour progression. Br J Cancer 2006, 95(10):1404–1409.

29. Rodriguez AC, Vahrenkamp JM, Berrett KC, Clark KA, Guillen KP, Scherer SD, Yang CH, Welm BE, Janat-Amsbury MM, Graves BJ et al: ETV4 Is Necessary for Estrogen Signaling and Growth in Endometrial Cancer Cells. Cancer research 2020, 80(6):1234–1245.

30. Bartha Á, Győrffy B: TNMplot.com: A Web Tool for the Comparison of Gene Expression in Normal, Tumor and Metastatic Tissues. Int J Mol Sci 2021, 22(5).

31. Fekete JT, Gyorffy B: ROCplot.org: Validating predictive biomarkers of chemotherapy/hormonal therapy/anti-HER2 therapy using transcriptomic data of 3,104 breast cancer patients. International journal of cancer Journal international du cancer 2019, 145(11):3140–3151.

32. Corbacioglu SK, Aksel G: Receiver operating characteristic curve analysis in diagnostic accuracy studies: A guide to interpreting the area under the curve value. Turk J Emerg Med 2023, 23(4):195–198.

33. Finlay-Schultz J, Jacobsen BM, Riley D, Paul KV, Turner S, Ferreira-Gonzalez A, Harrell JC, Kabos P, Sartorius CA: New generation breast cancer cell lines developed from patient-derived xenografts. Breast cancer research : BCR 2020, 22(1):68.

34. Menyhart O, Kothalawala WJ, Gyorffy B: A gene set enrichment analysis for cancer hallmarks. J Pharm Anal 2025, 15(5):101065.

35. Hilakivi-Clarke L, Warri A, Bouker KB, Zhang X, Cook KL, Jin L, Zwart A, Nguyen N, Hu R, Cruz MI et al: Effects of In Utero Exposure to Ethinyl Estradiol on Tamoxifen Resistance and Breast Cancer Recurrence in a Preclinical Model. Journal of the National Cancer Institute 2017, 109(1).

36. Wang H, Dai Y, Wang F: ETV4-mediated transcriptional activation of SLC12A5 exacerbates ferroptosis resistance and glucose metabolism reprogramming in breast cancer cells. Mol Med Rep 2024, 30(6).

37. Euhus D, Bu D, Xie XJ, Sarode V, Ashfaq R, Hunt K, Xia W, O’Shaughnessy J, Grant M, Arun B et al: Tamoxifen downregulates ets oncogene family members ETV4 and ETV5 in benign breast tissue: implications for durable risk reduction. Cancer Prev Res (Phila) 2011, 4(11):1852–1862.

38. Dumortier M, Ladam F, Damour I, Vacher S, Bièche I, Marchand N, de Launoit Y, Tulasne D, Chotteau-Lelièvre A: ETV4 transcription factor and MMP13 metalloprotease are interplaying actors of breast tumorigenesis. Breast Cancer Res 2018, 20(1):73.

39. Ladam F, Damour I, Dumont P, Kherrouche Z, de Launoit Y, Tulasne D, Chotteau-Lelievre A: Loss of a negative feedback loop involving pea3 and cyclin d2 is required for pea3-induced migration in transformed mammary epithelial cells. Mol Cancer Res 2013, 11(11):1412–1424.

40. Chen Y, Sumardika IW, Tomonobu N, Kinoshita R, Inoue Y, Iioka H, Mitsui Y, Saito K, Ruma IMW, Sato H et al: Critical role of the MCAM-ETV4 axis triggered by extracellular S100A8/A9 in breast cancer aggressiveness. Neoplasia 2019, 21(7):627–640.

41. Yuen HF, Chan YK, Grills C, McCrudden CM, Gunasekharan V, Shi Z, Wong AS, Lappin TR, Chan KW, Fennell DA et al: Polyomavirus enhancer activator 3 protein promotes breast cancer metastatic progression through Snail-induced epithelial-mesenchymal transition. J Pathol 2011, 224(1):78–89.

42. Clementz AG, Rogowski A, Pandya K, Miele L, Osipo C: NOTCH-1 and NOTCH-4 are novel gene targets of PEA3 in breast cancer: novel therapeutic implications. Breast cancer research : BCR 2011, 13(3):R63.

43. Giles ED, Wellberg EA: Preclinical Models to Study Obesity and Breast Cancer in Females: Considerations, Caveats, and Tools. J Mammary Gland Biol Neoplasia 2020.

44. Győrffy B: Survival analysis across the entire transcriptome identifies biomarkers with the highest prognostic power in breast cancer. Computational and Structural Biotechnology Journal 2021, 19:4101–4109.

45. Creighton CJ, Sada YH, Zhang Y, Tsimelzon A, Wong H, Dave B, Landis MD, Bear HD, Rodriguez A, Chang JC: A gene transcription signature of obesity in breast cancer. Breast cancer research and treatment 2012, 132(3):993–1000.

46. Park YH, Im SA, Park K, Wen J, Lee KH, Choi YL, Lee WC, Min A, Bonato V, Park S et al: Longitudinal multi-omics study of palbociclib resistance in HR-positive/HER2-negative metastatic breast cancer. Genome Med 2023, 15(1):55.

47. Rudolph MC, Wellberg EA, Lewis AS, Terrell KL, Merz AL, Maluf NK, Serkova NJ, Anderson SM: Thyroid Hormone Responsive Protein Spot14 Enhances Catalysis of Fatty Acid Synthase in Lactating Mammary Epithelium. Journal of lipid research 2014.

48. Meyerhardt JA, Irwin ML, Jones LW, Zhang S, Campbell N, Brown JC, Pollak M, Sorrentino A, Cartmel B, Harrigan M et al: Randomized Phase II Trial of Exercise, Metformin, or Both on Metabolic Biomarkers in Colorectal and Breast Cancer Survivors. JNCI Cancer Spectr 2020, 4(1):pkz096.

