## Supplemental Figures for "The Ets transcription factor ETV4 regulates FGF1-dependent proliferation and glycolysis in ER-positive breast cancer"

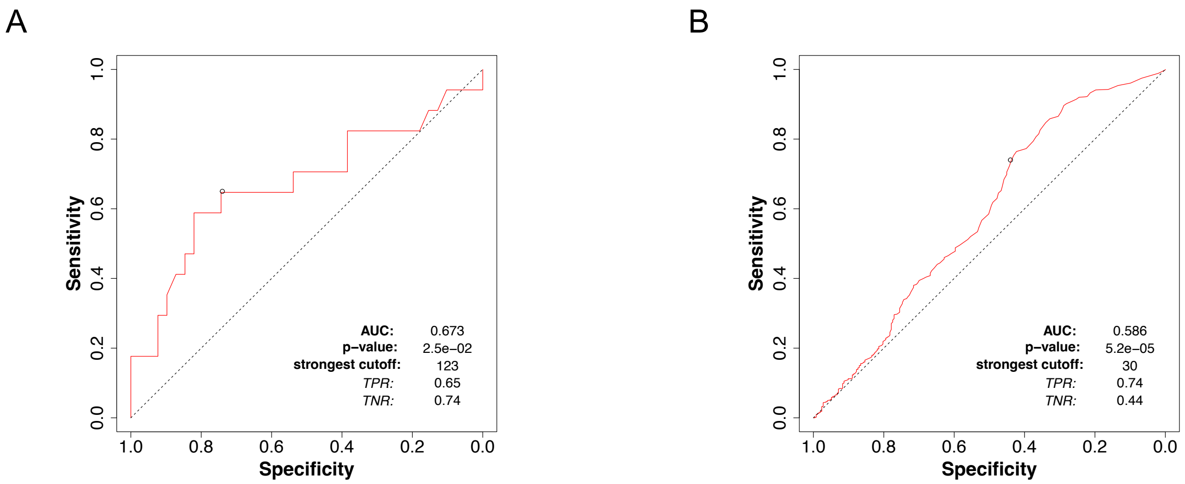


**Supplementary Figure 1. Utility of ETV4 as a prognostic biomarker.** Receiver operating characteristics (ROC) analysis showing the area under the curve of ETV4 expression in ER-positive breast tumors stratified by response to aromatase inhibition (a) or any chemotherapy (b). Data obtained from rocplot.com.

Supplemental Figure 2

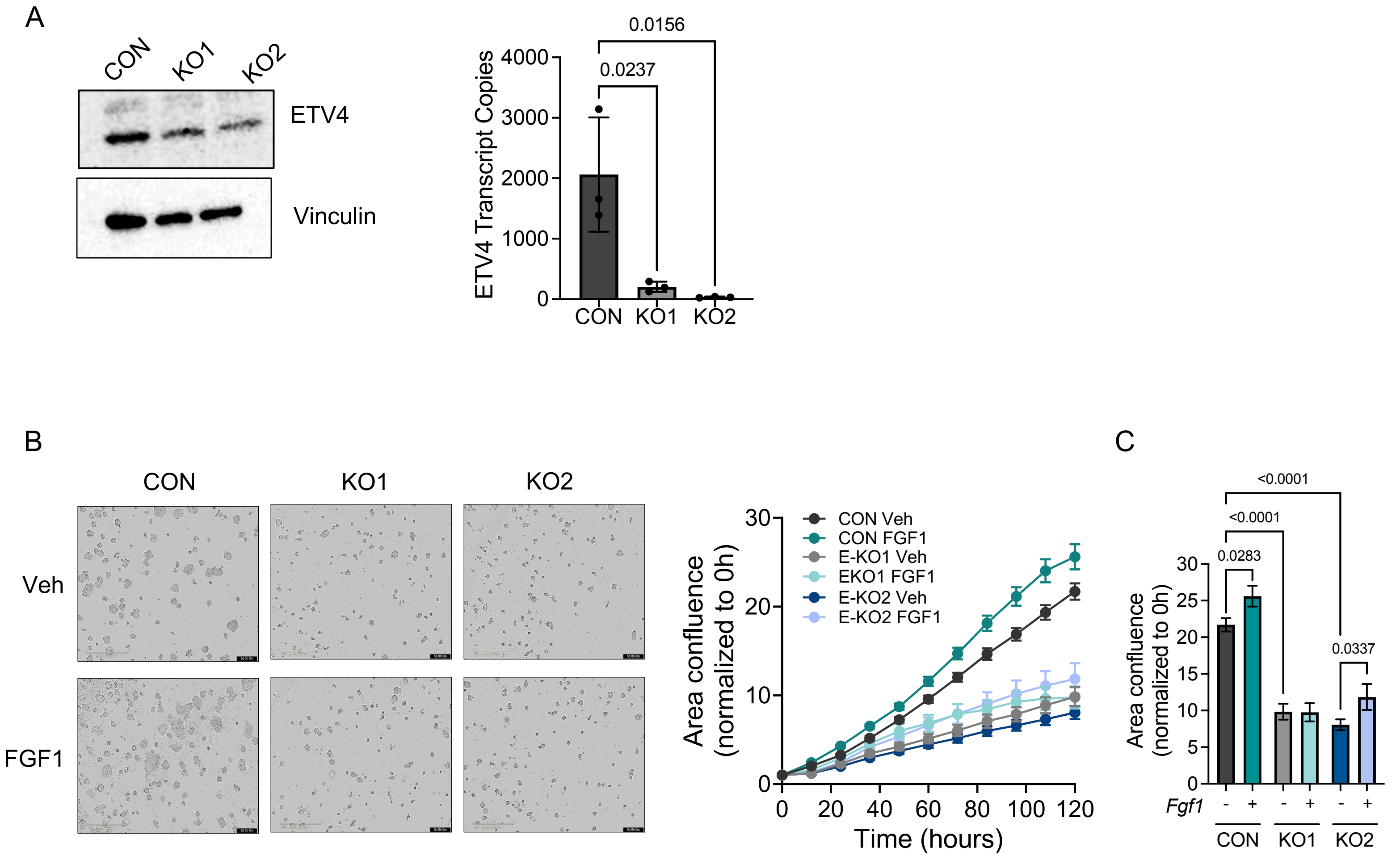


**Supplementary Figure 2. ETV4 loss suppresses FGF-1-mediated cell proliferation in UCD12 cells.** (A) Western blot analysis verifying ETV4 knockdown. (B) Q-PCR analysis of ETV4 expression. (C) Representative images of the final timepoint (left) and growth rates (right) of UCD12 control or ETV4-knockdown cells treated with vehicle or FGF1. (D) Area confluence relative to control vehicle time 0 of cells at the final timepoint following treatment. Two-way ANOVA testing for main effects of ETV4 knockdown or FGF1 treatment or interaction was performed. P-values indicate post-hoc multiple testing for specific differences between pre-defined comparisons.

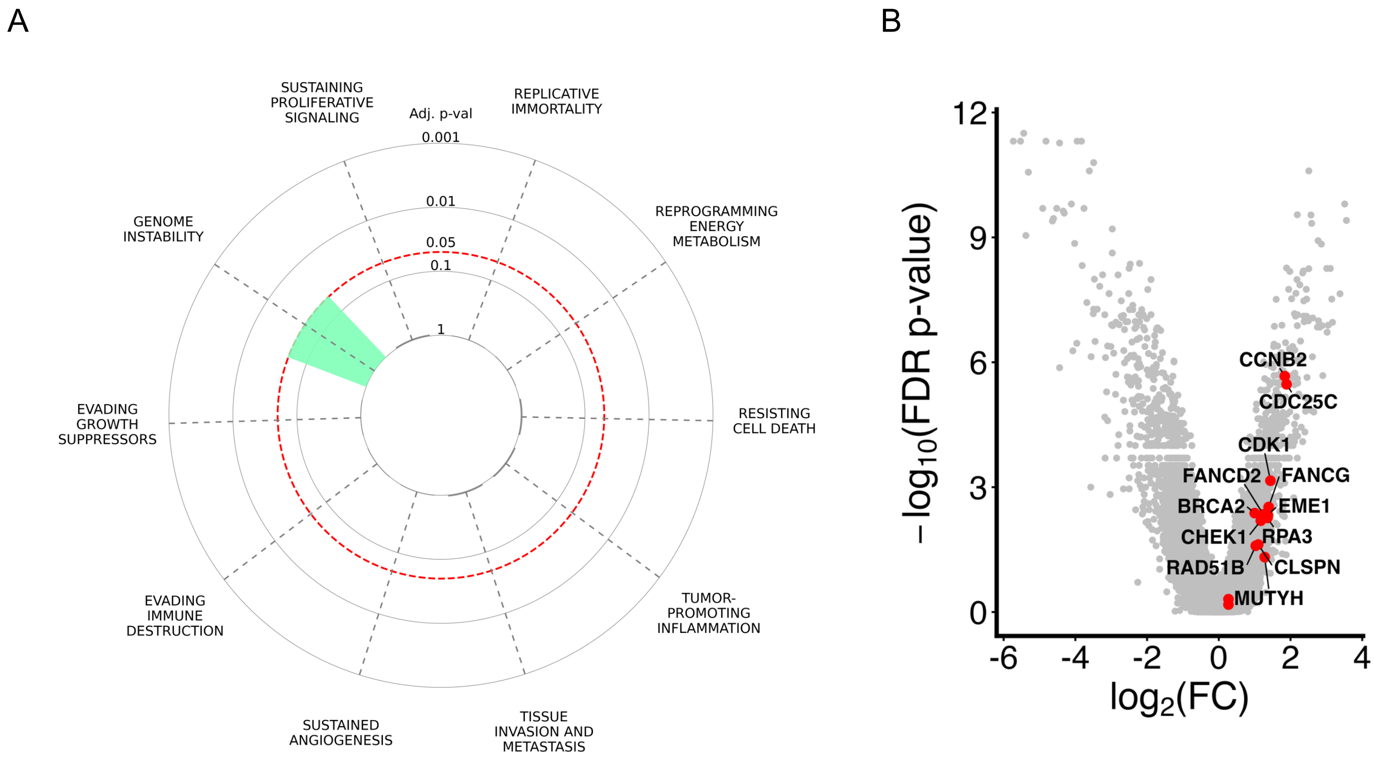


**Supplementary Figure 3: Cancer Hallmarks regulated by ETV4 loss.** (A) Hallmarks of Cancer enrichment plot illustrating pathways that are represented by differentially expressed genes ≥2-fold (adjusted p-value) in ETV4 knockdown compared with control vehicle in MCF7-TAMR cells. Bar height reflects −log10 adjusted p-value, with dashed circles indicating significance thresholds. (B) Volcano plot comparing ETV4 knockdown to control vehicle-treated MCF7-TAMR cells, highlighting upregulated genes involved in genome instability. Red points denote significantly upregulated genes.

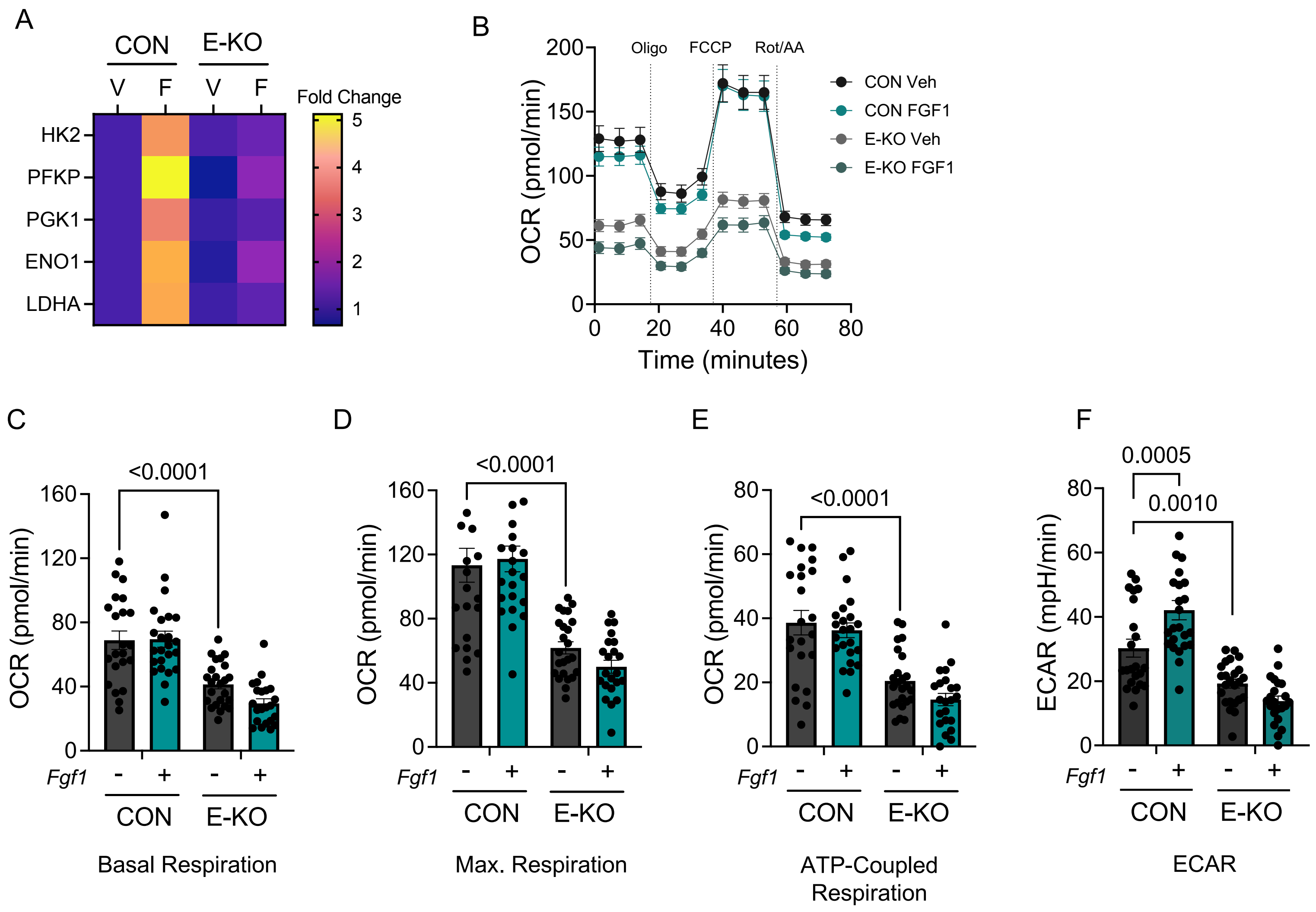


**Supplementary Figure 4: ETV4 loss alters glycolytic metabolism in UCD12 cells.** (A) Heatmap showing expression of glycolytic genes (HK2, PFKP, PGK1, ENO1, and LDHA) under vehicle and FGF1-treated conditions in UCD12 control and ETV4 knockdown cells. Data are expressed as fold change relative to the average of the vehicle for each gene, showing 3 replicates per group. (B) Seahorse metabolic flux analysis showing the kinetic graph of OCR in control and ETV4 knockdown UCD12 cells. (C-F) OCR parameters, including basal respiration (*c*), maximal respiration (*d*), ATP-production coupled respiration (*e*), and ECAR (*f*) in control and ETV4 knockdown UCD12 cells, upon vehicle and FGF1 stimulation. Data were analyzed using 2-way ANOVA.

### Supplemental Figure 5

A

Tumors Established

| *p=0.457* | Yes | No |
|---|---|---|
| CON | 7 | 7 |
| E-OE | 11 | 5 |

B

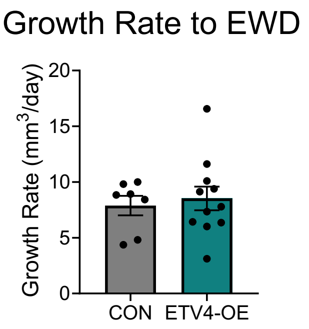


C

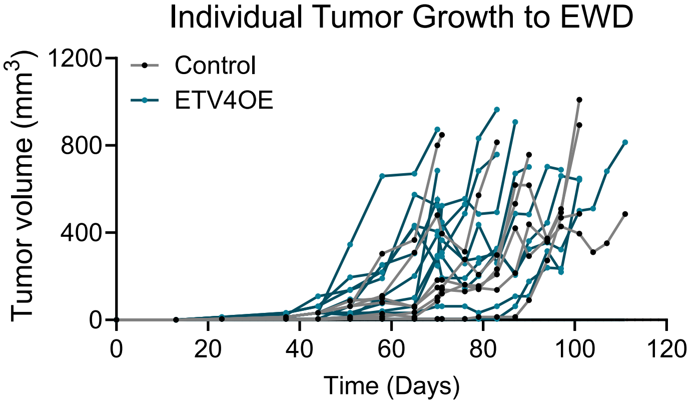


D

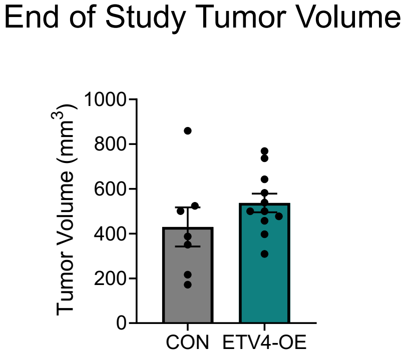


**Supplemental Figure 5. ETV4 overexpression does not impact tumor growth in lean mice.** (A) Tumor take rate in control and ETV4 overexpressing groups. Statistical significance was assessed using Fisher's exact test. (B) Tumor growth rate in mice until the time of EWD. (C) Individual tumor growth from the time of tumor injection to EWD in control and ETV4 overexpressing groups. (D) Tumor volume at the end of the study in control and ETV4 overexpressing groups.

A

Tumors Established

| *p=0.118* | Yes | No |
|---|---|---|
| CON | 12 | 2 |
| E-KO | 9 | 7 |

B

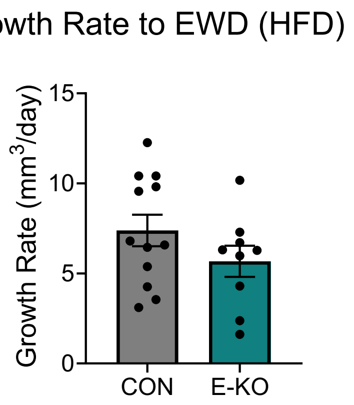


C

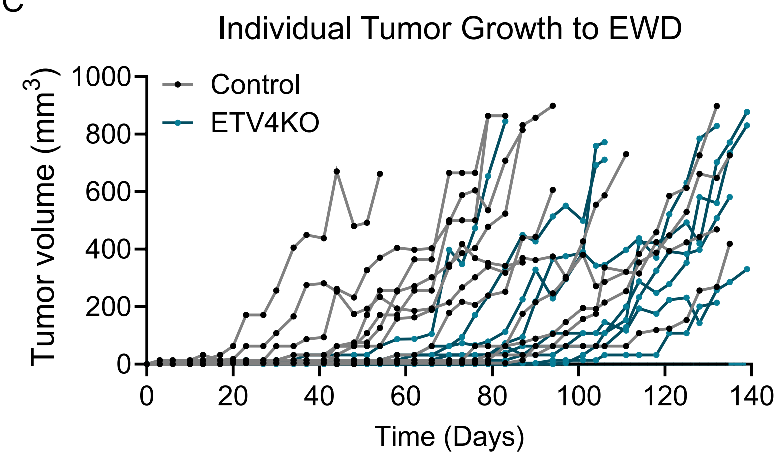


D

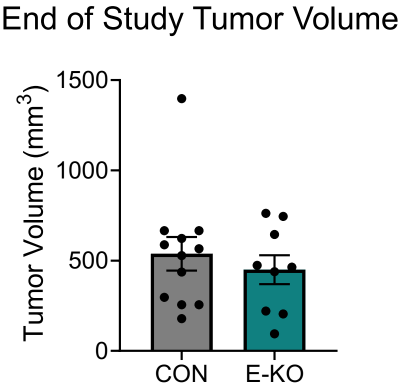


**Supplemental Figure 6. Loss of ETV4 in tumors from obese mice.** (A) Tumor take rate in control and ETV4 knockdown groups. Statistical significance was assessed using Fisher's exact test. (B) Tumor growth rate in mice until the time of EWD. (C) Individual tumor growth from the time of tumor injection to EWD in control and ETV4 knockdown groups. (D) Tumor volume at the end of the study in control and ETV4 knockdown groups. EKO = ETV4 knockdown.
